# Joint reconstruction of *cis*-regulatory interaction networks across multiple tissues using single-cell chromatin accessibility data

**DOI:** 10.1101/721290

**Authors:** Kangning Dong, Shihua Zhang

## Abstract

The rapid accumulation of single-cell chromatin accessibility data offers a unique opportunity to investigate common and specific regulatory mechanisms across different cell types. However, existing methods for *cis*-regulatory network reconstruction using single-cell chromatin accessibility data were only designed for cells belonging to one cell type, and resulting networks may be incomparable directly due to diverse cell numbers of different cell types. Here, we adopt a computational method to **j**ointly **r**econstruct *cis*-regulatory **i**nteraction **m**aps (JRIM) of multiple cell populations based on patterns of co-accessibility in single-cell data. We applied JRIM to explore common and specific regulatory interactions across multiple tissues from single-cell ATAC-seq dataset containing ~80,000 cells across 13 mouse tissues. Reconstructed common interactions among 13 tissues indeed relate to basic biological functions, and individual *cis*-regulatory network shows strong tissue specificity and functional relevance. More importantly, tissue-specific regulatory interactions are mediated by coordination of histone modifications and tissue related TFs, and many of them reveal novel regulatory mechanisms (e.g., a kidney-specific promoter-enhancer loop of clock-controlled gene *Gys2*).

## Introduction

*Cis*-regulatory elements (CREs) are a key class of regulatory DNA sequences and typically regulate the transcription of target genes by binding to transcription factors (TFs) (1,2). Lots of efforts have been made to characterize combinatorial patterns and systematically define CREs (3–6). Furthermore, interactions between CREs are vital components of genetic regulatory networks, controlling cell type-specific biological processes (7–9). However, linking CREs to their target genes is still a challenging problem due to their distal distance (in some cases hundreds of kilobases) and complicated regulatory mechanisms. Moreover, direct DNA contacts could be inferred using computational methods (e.g., CHiCAGO (10)) from data of the chromosome conformation capture (3C) technique and its variants such as ChIA-PET (11), Hi-C (12) and capture Hi-C (13).

But thus far these types of data are only available for a few cell types. As a result, several computational methods have been developed to estimate *cis*-regulatory DNA interactions using epigenetic data (14–18). For example, Corces *et al*. (14) proposed a computational strategy based on the correlation of transcriptional expression and chromatin accessibility to identify promoter-enhancer interactions. Cao *et al*. (15) adopted a random forest model to reconstruct enhancer-target networks using both histone modification data and chromatin accessibility data. However, the above methods were all designed to infer CREs interactions from bulk sequencing data. The limited number of samples per cell type make it hard to generate robust genome-wide *cis*-regulatory interaction maps.

Fortunately, the advent of single-cell ATAC-seq technology has enabled the genome-wide profiling of chromatin accessibility at single-cell resolution (19,20). Chromatin accessibility is a hallmark of active DNA regulatory elements, which delineates the *in vivo* availability of binding sites to TFs (21,22). These large-scale datasets (10^4^~10^5^ cells) provide a unique opportunity to detect cell type-specific CREs, assess the gene regulatory landscape, model the transcription factor regulatory grammar and reconstruct CREs interaction maps robustly using statistical models (23–26). For example, Cicero (26) employs a graphical lasso model to detect linkages between CREs based on co-accessibility patterns in single-cell data. Co-accessible DNA elements have been illustrated to be functionally related. Specifically, they exhibit physical proximity, are mediated by interacting TFs and undergo coordinated changes in histone modifications.

When working with single-cell data, identifying the differences between different cell types or the dynamic changes during cell development is a vital problem. Therefore, one might want to compare *cis*-regulatory interaction networks estimated from different cell types to investigate the common and specific regulatory patterns. The strategy adopting Cicero to reconstruct regulatory networks for each cell type respectively has several deficiencies: (1) resulting networks may be incomparable directly due to diverse numbers of cells as well as data sparsity in different cell types; (2) estimating separate networks does not exploit the similarity between multiple cell types, which will cause inaccurate results for identifying common patterns; (3) when we explore the cell type-specific *cis*-regulatory interactions, the uncorrelated technology noises of single-cell data from different cell types, which are caused by amplification process and batch effects (27), could lead to many false positives. Consequently, more accurate estimations are expected by considering multiple cell types simultaneously and alleviating uncorrelated technology noises.

To this end, we adopt a statistical method to **j**ointly **r**econstruct *cis*-regulatory **i**nteraction **m**aps (JRIM) of multiple cell populations based on patterns of co-accessibility in single-cell chromatin accessibility data. JRIM employs a group lasso penalty to encourage a similar pattern of sparsity across all the regulatory networks and alleviate uncorrelated technology noises. We applied JRIM to explore common and specific regulatory patterns across *cis*-regulatory interaction networks of different tissues using single-cell ATAC-seq dataset generated in (23), which profiled ~100,000 cells across 13 adult mouse tissues. Results show that common interactions of all tissues are significantly enriched in housekeeping gene regions, and common interactions of similar functional tissues are involved in regulations of tissue-shared biological processes. Furthermore, tissue-specific differential activity genes perform specific biological functions and have higher transcriptional expressions in the corresponding tissues. More interestingly, tissue-specific functional peaks are enriched for histone modification marks and motifs of TFs that are known to be important for tissue-specific functions. Last but not least, the reconstructed *cis*-regulatory interaction networks reveal distinct regulatory mechanisms of the sodium channel gene *Scn5a* and identify a novel kidney-specific promoter-enhancer loop of clock-controlled gene *Gys2*.

## MATERIALS AND METHODS

### Materials

Chromatin accessibility profiles at single-cell resolution in 13 adult mouse tissues have been generated using single-cell ATAC-seq technology in a recent study (23). We downloaded this single-cell ATAC-seq data from GSE111586. These 13 tissues include bone marrow, cerebellum, large intestine (including cecum and colon), heart, kidney, liver, lung, prefrontal cortex, small intestine (including duodenum, jejunum, and ileum), spleen, testes, thymus, and whole brain. To avoid batch effects, we only selected cells from one sample for each tissue. The processed single-cell ATAC-seq dataset contains ~80,000 cells and has called ~400,000 differentially accessible peaks using MACS2 (28) as described in (23). The total number of cells profiled per tissue ranges widely from 2,278 for cerebellum to 8,991 for heart **(Supplementary Figure S1A)**.

We collected available single-cell RNA-seq datasets for 11 tissues (except prefrontal cortex and whole brain). The single-cell RNA-seq data of cerebellum were collected from Dropviz database (29). Those of large intestine and heart were collected from Tabula Muris database (30). Those of other tissues were obtained from (31). We also collected available chromatin modification ChIP-seq datasets for 10 tissues, including H3K4me1, H3K4me3, H3K27ac and CTCF (except large intestine, prefrontal cortex and whole brain) from the ENCODE portal (https://www.encodeproject.org/) (32). And the genome-wide occupancy profiles of TBX3 in heart measured by ChIP-seq were collected from (33). In addition, we obtained mouse housekeeping gene set and 27 consistently expressed genes from (34) and collected tissue-specific differential expression genes of 12 tissues (except prefrontal cortex) from two mouse tissue-specific gene datasets (34,35).

### JRIM

Let ***X**^k^* be a *M* × *N_k_* matrix representing the binary accessibility values of the *k*-th tissue (*k* = 1,…, *K*), where 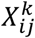 is 1 if one or more reads are observed to overlap the accessible peak *i* in cell *j* of tissue *k*, and 0 otherwise. JRIM takes the binary matrices of all *K* tissues as input **(Figure 1A)**, and consists of three main steps as follows.

**Figure 1.**
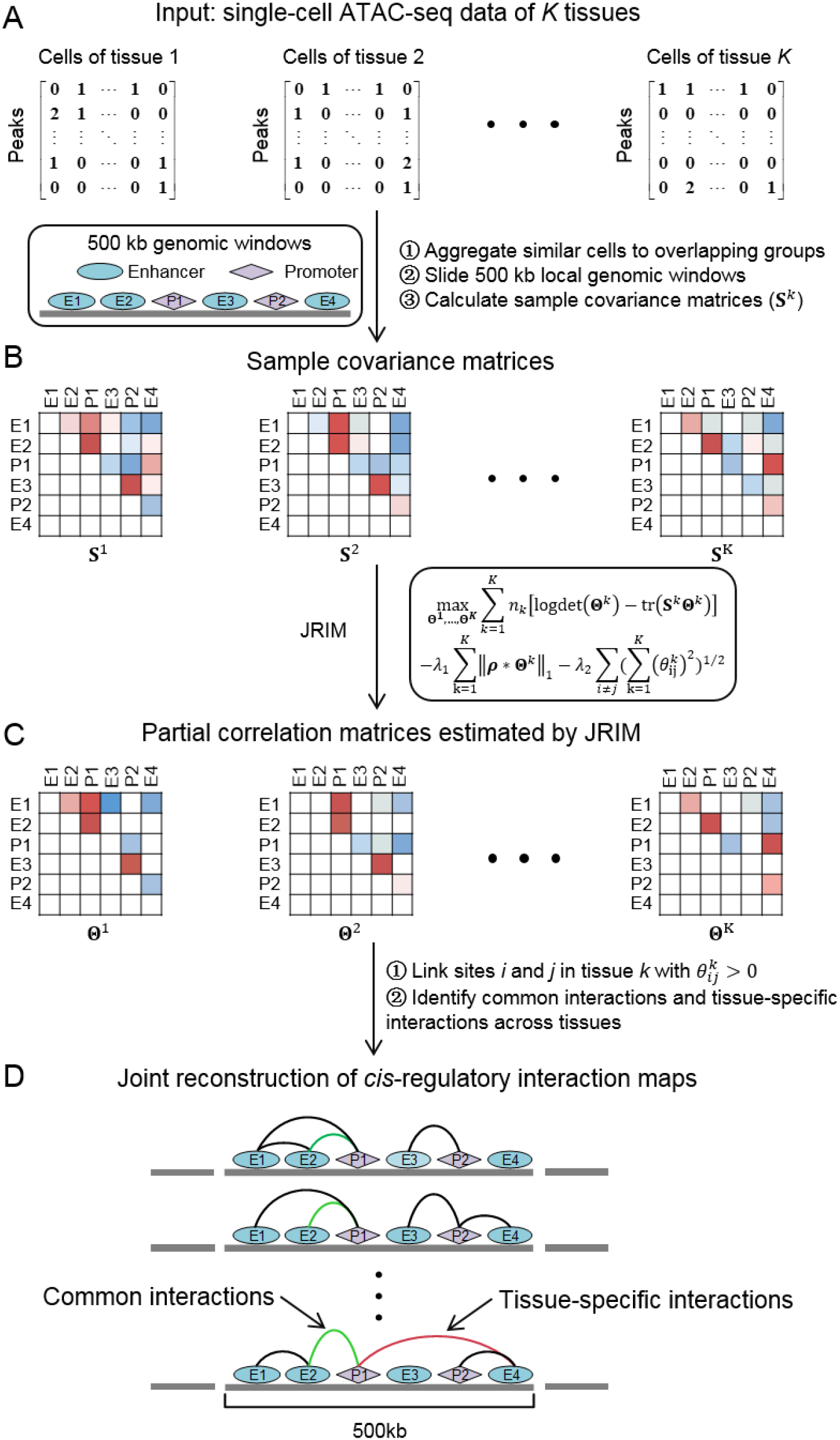
Illustration of the workflow of JRIM. **(A)** The input is single-cell chromatin accessibility data of multiple tissues. **(B)** JRIM aggregates similar cells to overlapping groups per tissue to overcome sparsity of single-cell data and calculates sample covariance matrices for local genomic windows. **(C)** JRIM jointly estimates local partial correlation matrices of *K* tissues and identifies co-accessible DNA element pairs, that is, *cis*-regulatory interactions. **(D)** Reconcile local regulatory interaction networks to achieve the reconstruction of genome-wide *cis*-regulatory interaction networks of all *K* tissues.

#### Step 1. Grouping similar cells per tissue

Sparsity of single-cell epigenetic data is intrinsic due to the limited DNA copy number, and the percentage of 0 in used single-cell ATAC-seq data is >98% **(Supplementary Figure S1B)**. The exceeding sparsity may cause several computational issues. For example, sample covariance matrix may be singular. To reduce the sparsity, control the technical variation and improve the stability of estimation, JRIM adopts a similar grouping procedure used as Cicero. This procedure can be regarded as a type of “bagging” (36).

JRIM first maps cells in each tissue into low dimensional space using a t-SNE map (37) and constructs a *k*-nearest neighbor graph via the *FNN* package (https://CRAN.R-project.org/package=FNN) **(Supplementary Figure S2)**. Then JRIM samples cells randomly and clusters their *k* nearest neighbors (default *k*=50). Binary accessibility profiles of cells in a group are aggregated into integer counts to construct the grouped matrix if a group does not overlap any existing one with more than 80% cells. This step will be performed for each tissue separately and generates grouped matrices **A**^1^, **A**^2^,…, **A**^*K*^ **(Figure 1B)**. Note that a cell will sometimes belong to more than one group. The different initializations of random selection will slightly influence the results.

#### Step 2: Adopting joint graphical lasso to compute co-accessibility scores between accessible sites in local genomic windows

JRIM jointly estimates sparse inverse covariance matrices, which encode partial correlations between variables, to capture the co-accessibility structure of accessible sites via a joint graphical lasso model (39) **(Figure 1C)**. Formally, JRIM employs the joint graphical lasso to maximize the following objective function:

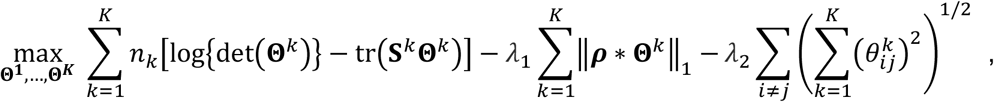

where **Θ**^*k*^ is the inverse covariance matrix of the aggregated count matrix **A**^*k*^, which captures the conditional dependence structure of accessible sites in tissue *k*, and **S**^*k*^ is the sample covariance matrix of **A**^*k*^. *n_k_* is the number of cell groups in tissue *k*, *λ*_1_ and *λ*_2_ are two non-negative tuning parameters, the matrix ***ρ*** encodes the distance penalty between accessible sites, and * denotes component-wise multiplication.

The first term of this formula is the maximum likelihood of estimating *K* graphical lasso models (40) respectively. To extract meaningful networks, the joint graphical lasso model expects only a small fraction of pairs to be partially correlated, i.e. to be non-zero in the inverse covariance matrices. Therefore, the second term is a *l*_1_-norm penalty to increase sparsity of the resulting inverse covariance matrices. The matrix ***ρ*** is determined by the following equation to induce distance penalty such that longdistance pairs have a relatively high penalty factor:

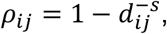

where *d_ij_* is the distance in the genome (in kilobases) between sites *i* and *j*, and *s* is a constant (*s* is set to 0.75 by default as suggested in Cicero).

The third term of this equation is the group lasso penalty (41) across all *K* inverse covariance matrices. This group lasso penalty encourages a similar pattern of sparsity across all the inverse covariance matrices—i.e. an interaction is encouraged to be *0*s in all the *K* estimated inverse covariance matrices if it only shows non-significant co-accessibility in a few tissues. This penalty helps to remove uncorrelated technology noises and exploit the similarity between multiple cell types.

Furthermore, as promoter capture Hi-C data suggested, >75% of three-dimensional promoter-based interactions occur within a 500kb distance (42). Therefore, JRIM pays more attention to local *cis*-regulatory interactions and estimates co-accessibility structure within overlapping 500kb genomic windows (windows are spaced by 250kb such that any accessible site is covered by two genomic windows).

#### Step 3: Constructing genome-wide *cis*-regulatory interaction maps

JRIM calculates partial correlations between local accessible sites for every 500kb genomic window. Then we will reconcile local interaction networks to construct genome-wide *cis*-regulatory interaction networks **(Figure 1D)**. Note that majority of DNA elements pairs are covered by two local co-accessibility networks and have two co-accessibility scores. In principle, these two co-accessibility scores are in the same direction (>95% in our experiment). If so, the mean score of these two pairs is assigned to the interaction in the resulting genome-wide *cis*-regulatory interaction networks. If not, such pairs are considered undetermined, and their co-accessibility values are set as zero. Finally, JRIM reconstructs genome-wide *cis*-regulatory networks where nodes are the accessible peaks, and edges link two co-accessible DNA elements which have co-accessibility scores above a user-defined threshold (set as 0 in this paper to avoid the impact of different distribution of co-accessibility scores across tissues).

### Selection of tuning parameters

The tuning parameters *λ*_1_, *λ*_2_ could control the sparsity and similarity of regulatory networks across all *K* tissues. However, both tuning parameters contribute to sparsity: *λ*_1_ encourages an individual network to be sparse and *λ*_2_ drives interaction intensity to be *0*s in all *K* networks. Therefore, as joint graphical lasso (39) recommended, we reparameterize *λ*_1_, *λ*_2_ as follows:

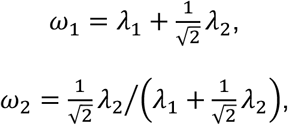

where *ω*_1_ and *ω*_2_ respectively reflect the levels of sparsity and similarity. To determine values of *ω*_1_ and *ω*_2_, JRIM first randomly selects 50 random 500kb genomic windows and calculates the minimum *ω*_1_ with a fixed *ω*_2_ (a relatively small value) such that non-zero entries in **Θ**^1^,…, **Θ**^K^ are less than 10% of all entries and no more than 5% long-distance pairs (pairs of sites at a distance greater than 250kb) are non-zeros. As for parameter *ω*_2_, we defined a similarity measurement as below:

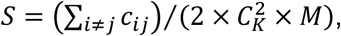

where *c_ij_* denotes the number of common non-zero pairs in tissues *i* and *j, K* is the number of tissues, 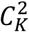 is the number of permutations, *M* is the mean number of non-zero pairs in all tissues. Then in the same 50 genomic windows, we fix *ω*_1_ as the mean of calculated values of *ω*_1_ and determine the minimum *ω*_2_ value such that measurement *S* is more than 0.6. Finally, the mean values of the calculated parameters are used for setting the penalties for each of 500kb genomic window sub-problem.

### Mapping *cis*-regulatory interactions to various genomic features

We obtained the genomic features including promoter regions (200bp upstream TSS and 100bp downstream TSS), 5’ UTR, 3’UTR, exons, introns and intergenic regions, which are located further than 1kb from any RefSeq gene, based on mm9 mouse reference genome from the UCSC Genome Browser (43). Firstly, peaks were categorized into the six feature categories if they overlap with any one genomic feature region. Then, *cis*-regulatory interactions were assigned into the same six feature categories according to accessible peaks they associated. Note that one peak may be mapped to more than one features and so do the interactions.

### Definition of tissue-specific differential activity genes (DAGs)

We depicted the activity of genes in a given tissue by the number of gene promoter related interactions in the reconstructed *cis*-regulatory interaction network. For gene *l*, we denoted the gene activity score vector as 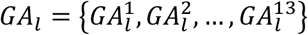, where 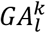 represents the number of promoter related interactions of gene *l* in the *k*th tissue. Then we normalized gene activity scores through dividing them by the total number of interacting co-accessible pairs in the corresponding tissue and calculated the z-scores of normalized gene activity scores 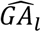:

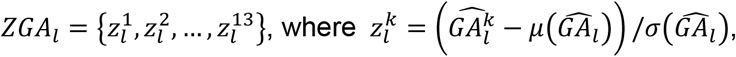

The gene *l* is considered as the *k*th tissue-specific differential activity one if 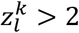.

### Exploring transcription and chromatin modification levels of tissue-specific DAGs

To test the transcription regulation functions of estimated *cis*-regulatory interactions, we investigated transcription levels and transcription specificity of DAGs using single-cell RNA-seq data. We first normalized the single-cell RNA-seq data of 11 tissues using the R package *Seurat* (44). For each tissue, genes were divided into two groups: ‘tissue-specific DAGs’ and ‘other genes’. We first calculated the mean expression of these two groups in their corresponding tissue. To characterize transcription specificity of DAGs, we also calculated the mean expression of ‘tissue-specific DAGs’ in other tissues.

We also investigated chromatin modification marks level around TSSs of DAGs to access the regulatory mechanisms. We calculated the mean signal of collected chromatin modification marks (H3K4me1, H3K4me3, H3K27ac and CTCF) in promoter region of all genes and normalized these signals to have the same mean value across all tissues. Then we grouped genes into two groups as in the analysis of gene expression data and compared their chromatin modification level.

### Gene ontology enrichment analysis

To assess the functional relevance of reconstructed interaction networks, we performed gene ontology (GO) enrichment analysis for common interaction related genes of similar functional tissues and tissue-specific differential activity genes via R package *clusterProfile* (45). Common interactions of similar functional tissues are defined as interactions that only detected in immune tissues (spleen, thymus, bone marrow and lung) or nervous tissues (cerebellum, prefrontal cortex and whole brain). We only selected genes whose promoters are associated with more than one common interaction of similar functional tissues to perform GO enrichment analysis. In the enrichment analysis of DAGs, we selected top 200 differential activity ones for each tissue (gene number of kidney, prefrontal cortex and large intestine are less than 200). The enrichment significance is corrected by Benjamini-Hochberg procedure.

### Motif analysis in tissue-specific interaction related regions

We defined the peaks relating to tissue-specific promoter-related interactions as ‘tissue-specific functional ones’. To investigate the regulatory mechanisms of tissue-specific interactions, we employed the HOMER software (http://homer.salk.edu/homer/) (46) to detect transcription factor motifs that are enriched in ‘tissue-specific functional peaks’ **(Table 1** and **Supplementary Table S1)**. The background sequences were generated from all accessible peaks. The brief description of TFs are obtained from UniProtKB database (https://www.uniprot.org/).

**Table 1.**
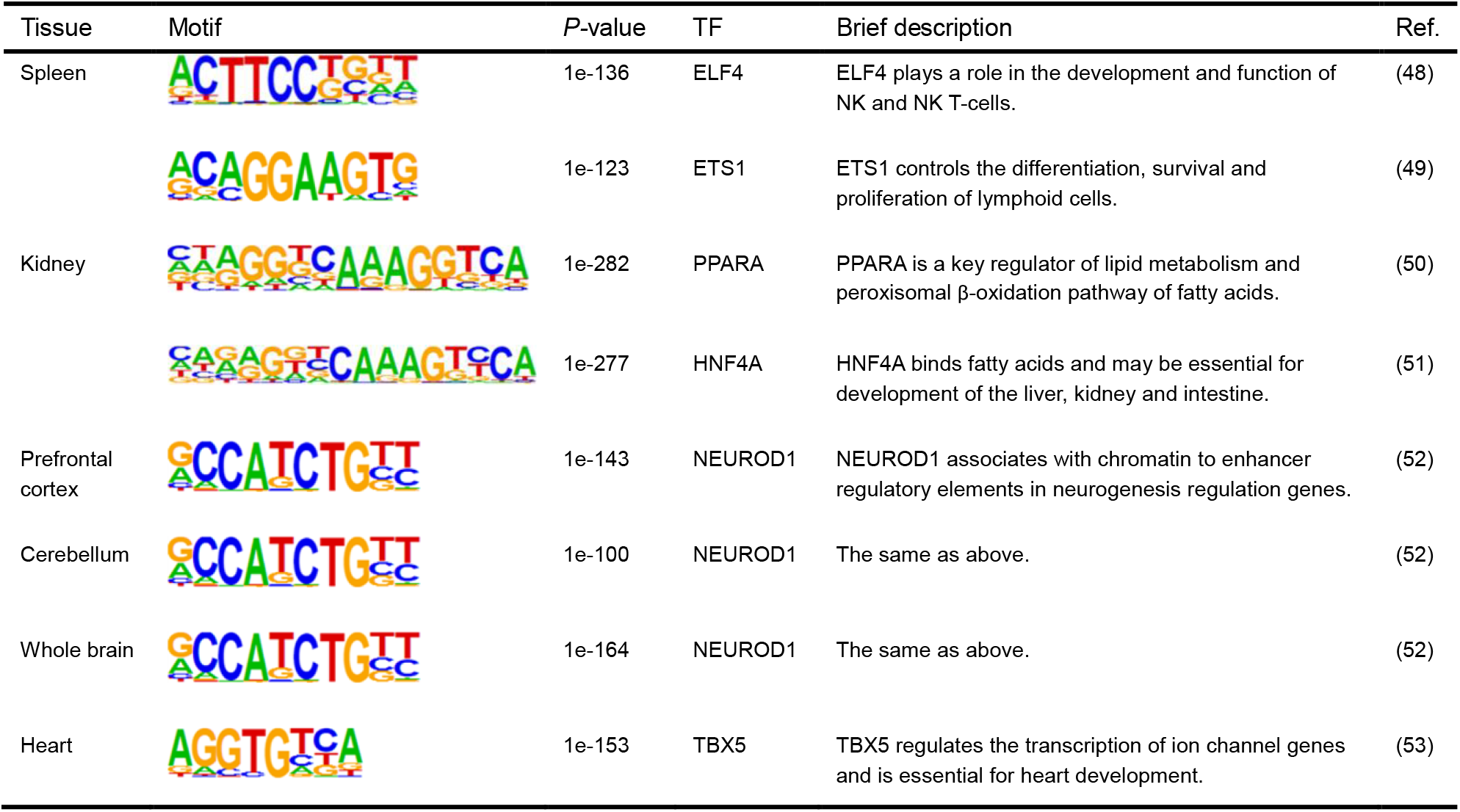
Examples of motifs enriched in the ‘tissue-specific functional peaks’. Examples are selected from the top two enriched motifs of each tissue.

| Tissue | Motif | *P*-value | TF | Brief description | Ref. |
| --- | --- | --- | --- | --- | --- |
| Spleen |  | 1e-136 | ELF4 | ELF4 plays a role in the development and function of NK and NK T-cells. | (48) |
|  |  | 1e-123 | ETS1 | ETS1 controls the differentiation, survival and proliferation of lymphoid cells. | (49) |
| Kidney | | 1e-282 | PPARA | PPARA is a key regulator of lipid metabolism and peroxisomal β-oxidation pathway of fatty acids. | (50) |
|  |  | 1e-277 | HNF4A | HNF4A binds fatty acids and may be essential for development of the liver, kidney and intestine. | (51) |
| Prefrontal cortex |  | 1e-143 | NEUROD1 | NEUROD1 associates with chromatin to enhancer regulatory elements in neurogenesis regulation genes. | (52) |
| Cerebellum |  | 1e-100 | NEUROD1 | The same as above. | (52) |
| Whole brain |  | 1e-164 | NEUROD1 | The same as above. | (52) |
| Heart |  | 1e-153 | TBX5 | TBX5 regulates the transcription of ion channel genes and is essential for heart development. | (53) |

## RESULTS

### JRIM jointly reconstructs comparable *cis*-regulatory networks of multiple tissues

We applied JRIM to jointly reconstruct *cis*-regulatory interaction maps of multiple mouse tissues from single-cell ATAC-seq data generated in (23). For the 436,206 accessible sites, we identified 2.28 million regulatory interactions in each tissue on average (ranging from 1.28 million in small intestine to 3.38 million in prefrontal cortex) **(Supplementary Figure S3A)**. Compared with Cicero (26), JRIM generates tissue-specific networks with relatively more comparable and balanced interaction**s (Supplementary Figure S3B)**. Proportions of tissue-specific interactions (only detected in one tissue) by JRIM have significantly decreased compared to those by Cicero, indicating JRIM characterize the shared interactions well among multiple tissues. The interactions frequency decreases with the increasing of genomic distance **(Supplementary Figure S4)**.

Note that the number of peak associated interactions is higher in transcription related genomic regions (especially in promoters and 5’ UTR), indicating their transcriptional regulation roles, and lower in intron as well as intergenic regions **(Figure 2A** and **Supplementary Figure S5)**. Moreover, we observed the same order when comparing conservation of interactions within different genomic regions, i.e. interactions associated with 5’ UTRs and promoters showed the highest conservation across tissues **(Figure 2B** and **Supplementary Figure S6)**, which is consistent with previous studies that DNA regulatory elements far from TSS exhibited a greater tissue specificity and wide dynamic range of activity (14,26,47).

**Figure 2.**
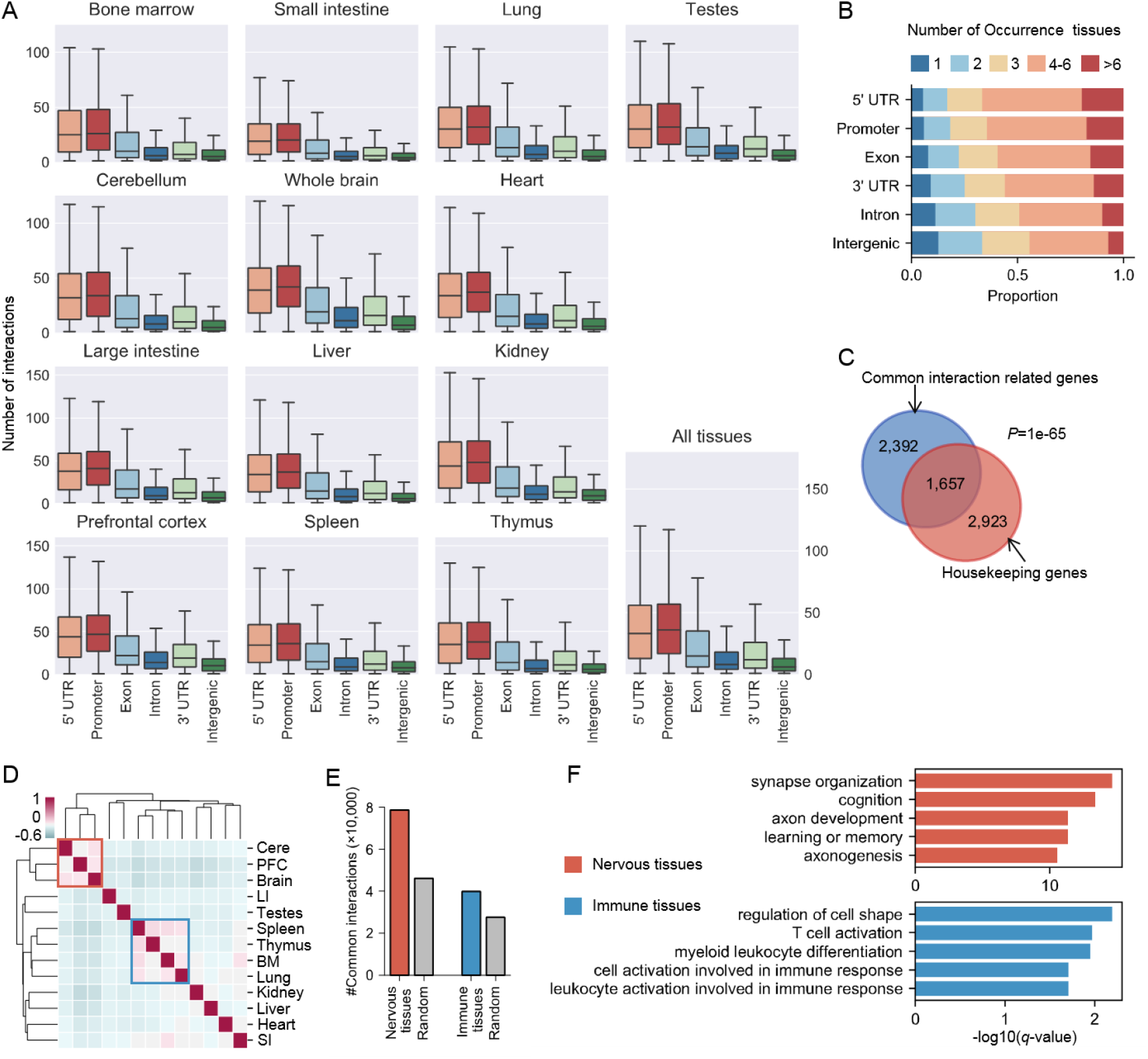
Basic characteristics of the reconstructed *cis*-regulatory interaction networks and the common interaction patterns across tissues. **(A)** Distribution of *cis*-regulatory interactions in six different genomic regions, including 5’ UTR, promoter, exon, intron, 3’ UTR and intergenic regions. **(B)** Distribution of conservation of genomic region related interactions. Color reflects the number of tissues detecting corresponding regulatory interactions. **(C)** Venn diagram of the overlapping between common interactions related genes and housekeeping gene set. The statistical significance was evaluated with Fisher’s exact test. **(D)** Hierarchical clustering of 13 tissues in terms of overlapping ratio of their *cis*-regulatory interactions. The two similar functional tissue clusters are marked by blue and red boxes. The values of heatmap represent Pearson’s correlation coefficients about the overlap ratio of *cis*-regulatory interactions. Cere: Cerebellum; PFC: prefrontal cortex; LI: large intestine; BM: bone marrow; SI: small intestine. **(E)** Comparison of the number of common interactions of similar functional tissue clusters (nervous tissues and immune tissues) and those of random selection tissues. **(F)** GO terms enrichment in common interaction related genes of similar functional tissues are associated with tissue sharing biological processes. Top 5 enriched GO terms of each cluster are shown.

### JRIM *cis*-regulatory networks reveal common interaction patterns among tissues

The comparable *cis*-regulatory interaction networks generated by JRIM enable direct identification of common or tissue-shared interactions among tissues. Note that the genes whose promoters are associated with common interactions of all 13 tissues are significantly enriched in housekeeping gene set (**Figure 2C;** *P*<1e-65, Fisher’s exact test), suggesting that these common interactions are involved in regulations of fundamental biological functions. We also observed that highly and constantly expression genes are regulated by more regulatory elements than remaining genes in all 13 tissues **(Supplementary Figure S7)**. Both observations suggest that the common patterns among all tissues are related to basic biological functions.

We observed that similar functional tissues are clustered together based on the overlapping ratio of their common interactions, including nervous tissues (cerebellum, prefrontal cortex and whole brain) and immune tissues (spleen, thymus, bone marrow and lung) **(Figure 2D)**. We also found that tissues within these two clusters have more common interactions than random selected ones **(Figure 2E)**. GO enrichment analysis of genes relating to common interactions of similar functional tissues (Materials and Methods) reveals distinct common biological processes of these tissues **(Figure 2F)**. For example, genes of common interactions for the nervous tissues are strongly enriched in ‘*synapse organization*’, ‘*cognition*’ and ‘*learning or memory*’ functional terms, and those of immune system are significantly enriched in ‘*T cell activation*’, ‘*myeloid leukocyte differentiation*’ and so on, indicating that the common interactions among tissues play key roles in regulations of their sharing biological functions.

### Reconstructed interaction networks show strong tissue specificity and functional relevance

Similar sparsity of *cis*-regulatory interaction networks enables quantification of the changes of gene activity in different tissues. Tissue-specific differential expression genes from two datasets (34, 35) both show higher activities in the corresponding tissue except tissue-specific genes of small intestine as well as the spleen-specific genes in TiSGeD dataset (**Figure 3A**; *p*<0.001 by one-sample Wilcoxon tests). Interestingly, the overlapping parts of two tissue-specific gene datasets tend to exhibit higher activity specificity compared with tissue-specific genes from an individual dataset. And the similar functional tissues (nervous tissues and immune tissues) are also clustered together in terms of gene activity scores **(Supplementary Figure S8)**. These results shed light on the underlying transcription regulation functions of *cis*-regulatory interactions and imply that tissue-specific gene expression is regulated by promoter-enhancer interactions.

**Figure 3.**
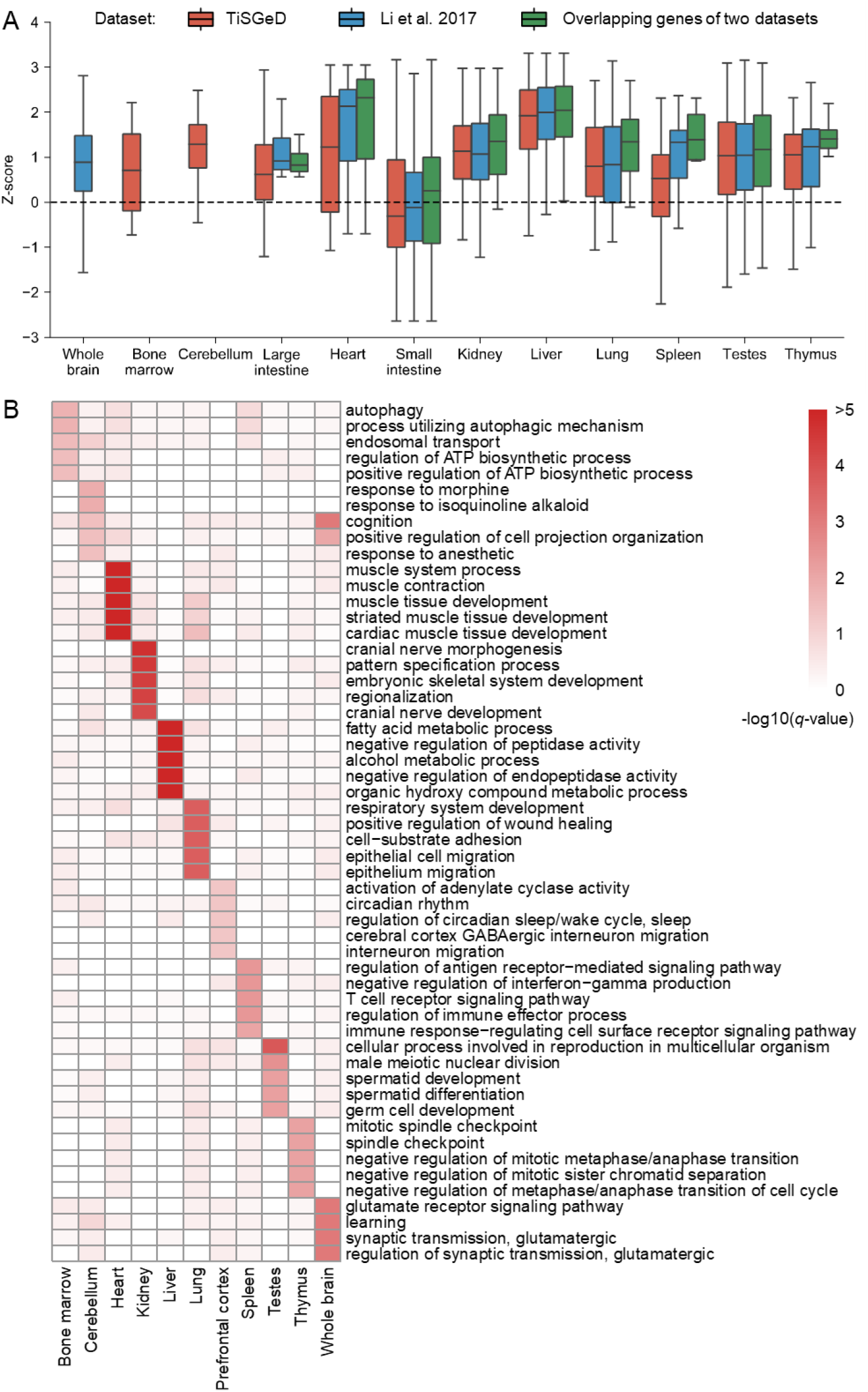
Tissue specificity and functional relevance of the reconstructed *cis*-regulatory interaction networks. **(A)** Boxplots of *z*-scores of two tissue-specific differential expression gene datasets and their overlapping genes. The z-scores are calculated by the number of promoter associated interactions in the reconstructed interaction networks. The dash line is the theoretical mean value of *z*-scores for randomly selected genes. **(B)** Differential activity genes for each tissue are enriched in tissue-specific biological processes. Top 5 enriched GO terms with *q*-value<0.05 for each tissue are selected. -log_10_(*q*-value) is used to plot this heatmap.

Functional enrichment analysis for tissue-specific differential activity genes (DAGs) (Materials and Methods) reveal that, except large intestine and small intestine, overrepresented GO terms were indeed relevant to the distinct tissue-specific biological functions **(Figure 3B)**. For example, tissue-specific DAGs of heart are enriched in cardiac muscle development related GO terms, such as ‘*cardiac muscle cell development*’, ‘*cardiac muscle tissue development*’ and so on. Those of testis are closely related to ‘*spermatid development*’ and ‘*germ cell development*’. Genes related to metabolism terms such as *‘fatty acid metabolic process’* and *‘alcohol metabolic process’* are strongly enriched in liver. As for nervous tissues, overrepresented GO terms are both related to neuron development or cognition functions, such as *‘cognition’* and *‘learning’* terms in both cerebellum and whole brain. These results highlight the underlying role of regulatory interactions in performing tissue-specific biological functions.

Next, we investigated the transcriptional behavior of tissue-specific DAGs and observed that tissue-specific DAGs have significantly higher transcriptional levels than other genes in all 11 tissues (**Figure 4A;** *p*<0.001, two sample Wilcoxon tests), highlighting the potential tissue-specific functions of DAGs. Moreover, tissue-specific DAGs from all tissues except those of small intestine exhibit strong transcriptional specificity (**Figure 4A**; *p*<0.01 for thymus and *p*<0.001 for others, two sample Wilcoxon tests). Thus, the reconstructed promoter-enhancer interactions are involved in gene activation to perform tissue-specific biological functions.

**Figure 4.**
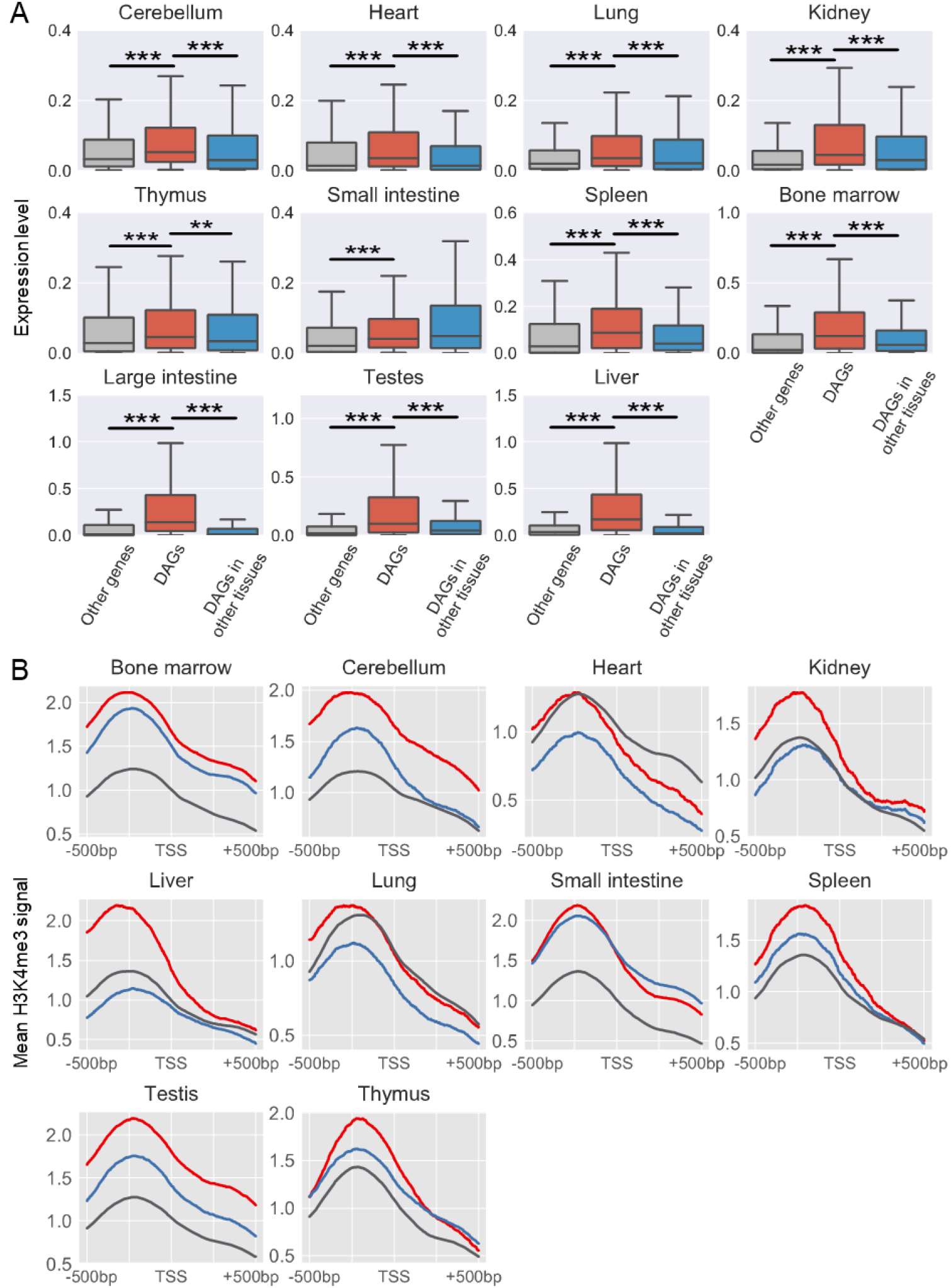
Transcription and histone modification levels of tissue-specific differential activity genes. **(A)** Boxplots of transcription levels of differential activity genes (DAGs) and other genes in the single-cell RNA-seq dataset. The middle red box represents RNA expression level of DAGs in the corresponding tissues. The left gray box represents expressions of other genes. The right blue box represents these DAGs expression in other tissues. Each middle box was compared with both left and right boxes using two-sample Wilcoxon tests. ** and *** indicate 0.001<*P*<0.01 and *P*<0.001 respectively. **(B)** Enrichment analysis of H3K4me3 mark around TSSs of DAGs in ten tissues. The signals around DAGs TSSs and TSSs of remaining genes in the corresponding tissue are labeled as red and gray colors, respectively. The blue line is the mean signals around TSSs of DAGs in other tissues.

We also found that chromatin modification marks around TSSs of tissue-specific DAGs in the corresponding tissue exhibit strong difference compared with those in other tissues **(Figure 4B** and **Supplementary Figure S9)**. Specifically, the active promoter mark H3K4me3 is strongly enriched in regions around TSSs of DAGs in most tissues (except heart and lung) **(Figure 4B)**. And H3K4me3 also shows tissue-specific enrichment around TSSs of tissue-specific DAGs, except those of small intestine **(Figure 4B)**, supporting the activity specificity of DAGs identified in the reconstructed networks by JRIM. Meanwhile, the signals of the architectural protein CTCF around TSSs of DAGs are also significantly higher in the corresponding tissue than those in other ones **(Supplementary Figure S9A)**, indicating that the promoter of tissue-specific DAGs tend to be related to chromatin looping in the given tissues. Moreover, the H3K27ac mark, known to be associated with the activation of transcription, is also strongly enriched in promoter regions of tissue-specific DAGs **(Supplementary Figure S9B)**. These observations indicate that epigenetic features may contribute to the regulation functions of promoter-enhancer interactions.

### Tissue-specific *cis*-regulatory interactions are mediated by coordination of tissue related TFs and histone modifications

JRIM enables the identification of tissue-specific interactions and investigation of tissue-specific regulatory mechanisms. The tissue-specific functional peaks associated with tissue-specific interactions are enriched for motifs of TFs that are functionally associated with tissue functions (Materials and Methods; **Table 1** and **Supplementary Table S1**). For instance, motifs of ELF4 and ETS1 were significantly overrepresented in spleen-specific functional peaks (*P*=1e-136 and *P*=1e-123, respectively). Note that ELF4 plays a role in development of NK and NK T-cells (48), and ETS1 controls the differentiation, survival and proliferation of lymphoid cells (49). In other examples, motifs of PPARA and HNF4A are significantly enriched in kidney-specific functional peaks (*P*=1e-282 and *P*=1e-277, respectively). PPARA has been reported to be a key regulator of lipid metabolism (50), and HNF4A is essential for development of kidney (51). Moreover, NEUROD1 was shown to be associated with neurogenesis (52), and motifs of NEUROD1 are highly enriched in prefrontal cortex, cerebellum and whole brain (*P*=1e-143, *P*=1e-104 and *P*=1e-164, respectively). We also investigated histone modification markers in tissue-specific functional peaks. Indeed, the enhancer marks (H3K27ac and H3K4me1) are significantly enriched in tissue-specific functional peaks **(Figure 5A** and **Supplementary Figure S10A**), highlighting the potential regulation function of these regions. Moreover, the CTCF marker is also enriched in tissue-specific functional peaks (**Supplementary Figure S10B**), which might be related with tissue-specific chromatin looping. Overall, the reconstructed *cis*-regulatory networks by JRIM could reveal tissue-specific interactions, which are mediated by coordination of tissue related TFs and histone modifications.

**Figure 5.**
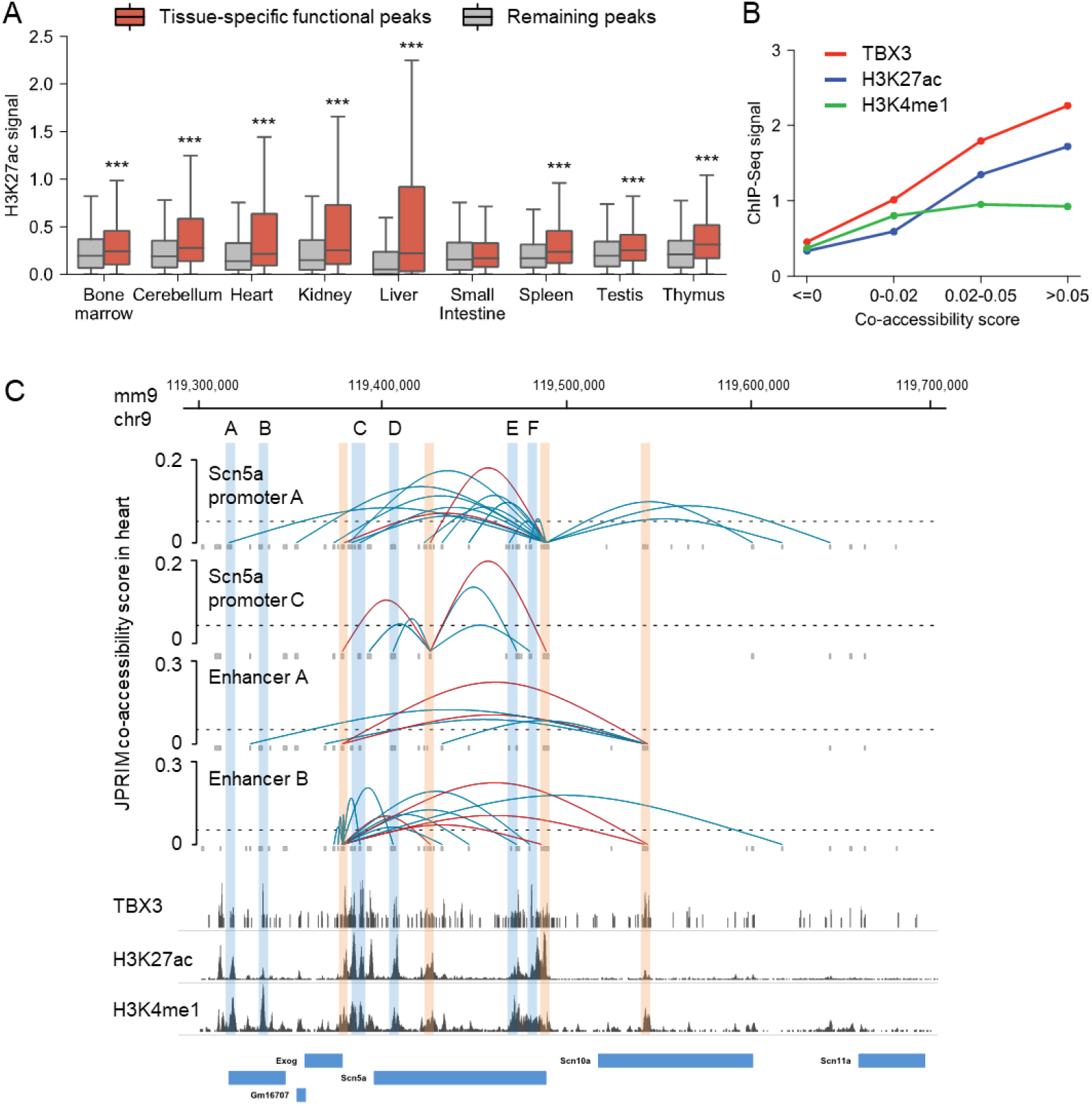
Illustration of the distinct regulatory mechanisms of *Scn5a*. **(A)** Boxplots of the H3K27ac signal of tissue-specific functional peaks compared to those of other peaks. **(B)** The occupancy profiles of TBX3, H3K4me1 signal and H3K27ac signal of *Scn5a* promoter-related peaks (co-accessibility score > 0) and other peaks. **(C)** The reconstructed *cis*-regulatory interaction network around the *Scn10a-Scn5a* locus (chr9: 119,300,000-119,700,000) in heart. The regions of *Scn5a* promoters and enhancer A, B are labeled with red boxes. Blue boxes indicate locations of TBX3 enriched regions A-F. The bottom is ChIP-seq data around *Scn5a* gene. The image is drawn on basis of WashU epigenome browser with RefSeq gene annotations.

### JRIM *cis*-regulatory networks reveal distinct regulatory mechanisms of the sodium channel gene Scn5a

The *cis*-regulatory interaction network of heart around *Scn10a-Scn5a* locus provides a strong illustration of cooperative regulatory roles about TF binding and histone modification. Motifs of TBX5 are significantly enriched in heart-specific functional peaks (**Table 1**, *P*=1e-153). Note TBX5 is known to activate ion channel genes and is essential for heart development (53). In addition, TBX3 is also critical for the development of heart, which competes with TBX5 for the same binding sites, but represses the expression of ion channels genes, such as the sodium channel gene *Scn5a* (54). In the reconstructed *cis*-regulatory interaction network of heart, the occupancy profile of TBX3 shows distinctly higher enrichment in the peaks linked to *Scn5a* promoter A (accessible peak: chr9: 119,487,317-119,490,317) **(Figure 5B)**, implying that *Scn5a*-related regulatory interactions in heart are mediated by bindings of TBX3 and TBX5. Meanwhile, the enhancer marks H3K27ac and H3K4me1 are also strongly enriched in *Scn5a* promoter-related regions **(Figure 5B)**. More interestingly, with the increase of co-accessibility scores, the signals of TBX3 and H3K27ac also enhance a lot, but the signals of H3K4me1 are relatively stable. H3K27ac has been reported to be used to find active enhancers by subtracting H3K4me1 (55). This suggests that some histone modifications contribute to the binding of TFs, and the binding of TFs in active enhancer region have a greater impact on gene regulations.

The reconstructed *cis*-regulatory interactions related to *Scn5a* promoters are associated with TBX3 enriched regions **(Figure 5C)**. Two of these TBX3 enriched regions have been well-documented to modulate the expression of *Scn5a* (33,56,57) (a region located in Scn10a gene region labeled as enhancer A: chr9:119,540,800– 119,544,032 and an intergenic region labeled as enhancer B: chr9:119,378,051–119,379,479). Specifically, Boogaard *et al*. (57) has leveraged high-resolution 4C-seq to investigate the chromosome structure between enhancer A, enhancer B and promoters of *Scn5a*, finding that these two enhancers and *Scn5a* promoters are in close contact and form a complex. They also engineered deletions of enhancer A and B and found that removal of these two enhancers significantly abrogates the ventricular conduction system and the expression of *Scn5a*, indicating the necessity of enhancer A and B for *in vivo Scn5a* expression regulation.

The validated interactions between enhancer B and *Scn5a* promoters were identified with high co-accessibility scores in the *cis*-regulatory network of heart, supporting the transcription regulation role of enhancer B **(Figure 5C)**. However, as for enhancer A, we only observed the regulatory contacts between enhancer A and enhancer B in heart **(Figure 5C)**. And the H3K4me1 signal in enhancer A region is relatively high, but the H3K27ac signal is low. We guessed that although the physical proximity between enhancer A and *Scn5a* promoters and the bindings of TBX3 and TBX5 in enhancer A region, enhancer A regulates the expression of *Scn5a* through indirect interactions (enhancer A-enhancer B-promoters of *Scn5a*) due to its low enhancer activity. In short, the regulatory network of heart around *Scn10a*-*Scn5a* locus reveals distinct regulatory mechanisms of *Scn5a*: regions enriched in both TBX3 bindings and H3K27ac are likely to directly regulate the expression of *Scn5a* (region A, C-F and enhancer B in **Figure 5C**) and regions only enriched in TBX3 bindings tend to not interact with *Scn5a* promoters (region B and enhancer A in **Figure 5C**). Moreover, the regulatory mechanisms of enhancer A demonstrate that physical proximity does not definitively mean regulatory interaction. Therefore, reconstructing *cis*-regulatory networks using chromatin accessibility data could help to better understand regulatory mechanisms and guide downstream analysis (see another example in **Supplementary Figure S11**).

### JRIM identifies a novel long-range kidney-specific promoter–enhancer loop of clock-controlled gene *Gys2*

We further investigated the regulatory interaction networks around Glycogen Synthase 2 (*Gys2*) gene, which is a liver-specific clock-controlled gene, catalyzing the rate-limiting step in hepatic glycogen synthesis (58). The transcript of *Gys2* is rhythmically expressed in liver during a day. In contrast, the accumulation of *Gys2* mRNA is constant and low in kidney (59). We compared the *Gys2* promoter related interactions in all 13 tissues and found a liver-specific contact with a region of 12kb to 21kb downstream from *Gys2* TSS **(Figure 6A)**. Two of three peaks located in this intragenic region exhibit higher co-accessibility with *Gys2* promoter in liver comparing to those in kidney **(Figure 6B)**. A recent study investigated the clock-dependent chromatin topology of *Gys2* in kidney and liver (59). They measured the interaction frequencies of *Gys2* promoter at different time points in liver and kidney using 4C-seq technology and found that the promoter of *Gys2* contacts with an intragenic region, 21 kb downstream from the TSS of *Gys2*, more frequently at the day versus at the night only in liver **(Figure 6C** and **Supplementary Figure S12A**). To verify this liver-specific promoter-enhancer loop, they cultured clock-deficient *Bmal1* knockout mice through hybridization and found that rhythms of this liver-specific interaction are abolished **(Supplementary Figure S12B)**. This putative liver-specific promoter-enhancer loop is highly consistent with our observation in the reconstructed *cis*-regulatory networks. Moreover, comparing the *cis*-regulatory interaction networks of kidney and liver around *Gys2*, JRIM detects a long-range interaction within a region of 280kb to 320kb downstream from *Gys2* TSS only in kidney **(Figure 6A)**. This region is suggested to be involved in a kidney-specific promoter-enhancer loop. As expected, the comparison of liver and kidney 4C-seq data shown significant increased contact frequency with the *Gys2* promoter within this region in kidney **(Figure 6C)**. We employed *FourCSeq* to estimate the spatial interactions of *Gys2* promoter in liver and kidney using 4C-seq data (60), finding a long-range kidney-specific chromatin loop within this region **(Figure 6D** and **Supplementary Figure S13**). Furthermore, we compared the histone modification marks in this region and observed that the enhancer marks (H3K4me1, H3K27ac) of peaks within this region in kidney are far greater than those in liver **(Figure 6E)**. The physical proximity and the higher histone modification level in kidney indicated that this region modulates expression of *Gys2* through a long-range promoter-enhancer loop in kidney.

**Figure 6.**
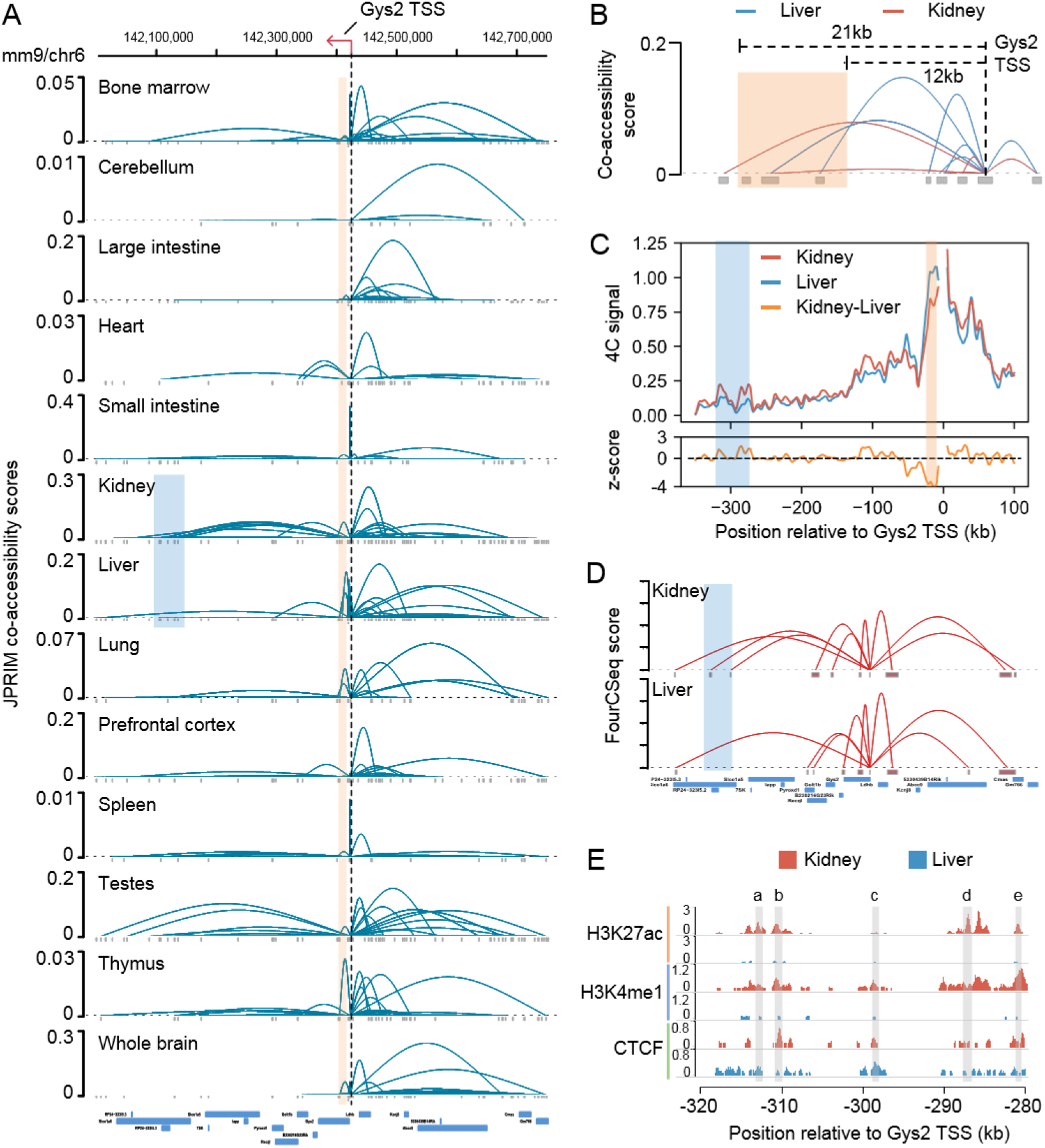
Illustration of two tissue-specific enhancer-promoter interactions revealed by the reconstructed networks. **(A)** Visualization of *Gys2* promoter related interactions within region (chr6:142,000,000-142,750,000) in all 13 mouse tissues. The black dashed line indicates the location of *Gys2* TSS. The red box indicates the genomic location of validated liver-specific enhancers. The blue box indicates a region located from 280kb to 320kb downstream of *Gys2* promoter. **(B)** Comparison of regulatory networks of kidney and liver in the region from 25kb downstream to 10kb upstream of *Gys2* promoter. The red box indicates the same genomic location as in (A). **(C)** The 4C-seq signal in a 450kb genomic region surrounding *Gys2* in kidney and liver. The 4C-seq profiles depict the interaction frequencies of *Gys2* promoter. The red box and blue box are the same as in (A). **(D)** Spatial interaction networks around *Gys2* TSS in kidney and liver estimated by *FourCSeq* using 4C-seq data. The blue box indicates the same region as in (A). **(E)** Signals of three chromatin modification marks across the region located from 280kb to 320kb downstream of *Gys2* promoter. Grey boxes indicate the location of accessible peaks a–e.

## DISCUSSION

Exploring tissue-specific *cis*-regulatory interaction patterns across multiple cell types could help to investigate cell type-specific regulation. Here, we adopt JRIM to jointly reconstruct *cis*-regulatory interaction maps of multiple cell populations from single-cell chromatin accessibility data. JRIM controls the sparsity of resulting *cis*-regulatory interaction networks simultaneously and alleviates uncorrelated technological noises across different cell populations. These *cis*-regulatory interaction networks could advance the understanding of regulatory mechanisms and guide downstream analysis such as CRISPR-mediated (epi)-genome editing.

Although we focused on tissue level and did not emphasize the cell heterogeneity within tissues in this study, the benefits of considering cell heterogeneity can be demonstrated through the poor functional relevance of small intestine-specific differential activity genes. Unlike other tissues, the profiled cells of small intestine were dissected from different regions including duodenum, jejunum, and ileum. The detected co-accessibility patterns (number of interactions) in small intestine are much fewer than other tissues **(Supplementary Figure S3A)**, which is probably caused by cell heterogeneity. JRIM could be applied to chromatin accessibility data with respect to cell populations defined ahead to obtain accurate regulatory networks.

Comparing to Cicero, JRIM exploits the similarity between tissues and generates *cis*-regulatory networks that can be compared directly. In a simple experiment, we found that although the two scaling parameters of JRIM *λ*_1_, *λ*_2_ contribute to the sparsity of networks both, the combined parameters *ω*_1_, *ω*_2_ separately control the sparsity level and similarity among reconstructed networks **(Supplementary Figure S14)**. Especially when fix *ω*_1_, enlarging *ω*_2_ brings almost no change in sparsity, but the similarity among tissues is increasing. We regarded it as an important merit relative to Cicero, and JRIM could generate biologically plausible results based on the data-driven parameter selection process (Materials and Methods).

Currently, most tissue-specific genes are identified on the basis of differential expression or tissue diseases related coding-region mutations. In this paper, we identified tissue-specific differential activity gene sets based on the changes of promoter associated interactions across tissues. We have shown that tissue-specific differential activity genes tend to be highly expressed in corresponding tissues **(Figure 4A)**. Moreover, the mutations within some DAG regions have been reported to be associated with tissue-related complex diseases on mouse or human. Taking heart as an example, the mouse embryos harboring a cardiomyocyte-restricted mutation in the *Myocd* gene exhibit myocardial hypoplasia and defective atrial (61), recessive *Ttn* truncating mutations in human are known to be associated with core myopathies (62), and *Myh6* mutations affect myofibril formation and are associated with congenital heart defects (63). The methodologies developed here for identifying tissue-specific genes may provide new biomarkers for the illustration of mutation and disease-related regulatory mechanisms.

In this work, we mainly focused on the direct transcriptional regulations, i.e. protein coding gene related promoter-enhancer interactions. However, as shown in the case study of *Scn5a*, indirect interactions also play a critical role in regulations of gene expression. Moreover, long intergenic noncoding RNAs (lincRNAs) are known for important regulators of gene expression (64), and aberrant epigenetic patterns in lincRNAs are might associated with cancer development (65). Therefore, characterizing the regulatory functions of indirect interactions and non-coding regions will be an intriguing problem. Deciphering modular structure of the reconstructed *cis*-regulatory networks can be used to distinguish biological function units and shed light on understanding of non-coding genes as well as non-coding mutations. Furthermore, a very interesting question in single-cell biology is modeling the dynamic changes during cell development. JRIM could detect dynamic changes of *cis*-regulatory interaction networks during cell development by analyzing time-series single-cell ATAC-seq data (24,66,67) or coupling with ‘pseudotime’ algorithms such as Monocle 2 (68).

## AVAILABILITY

The package of JRIM is available at http://page.amss.ac.cn/shihua.zhang/software.html. All datasets in this manuscript are public datasets.

## SUPPLEMENTARY DATA

Supplementary data are available upon request.

## REFERENCES

1. Wittkopp, P.J. and Kalay, G. (2012) Cis-regulatory elements: molecular mechanisms and evolutionary processes underlying divergence. Nat. Rev. Genet., 13, 59–69.

2. Zhang, S., Tian, D., Tran, N.H., Choi, K.P. and Zhang, L. (2014) Profiling the transcription factor regulatory networks of human cell types. Nucleic Acids Res., 42, 12380–12387.

3. Ernst, J. and Kellis, M. (2012) ChromHMM: automating chromatin-state discovery and characterization. Nat. Methods, 9, 215–216.

4. Chen, C., Zhang, S. and Zhang, X.-S. (2013) Discovery of cell-type specific regulatory elements in the human genome using differential chromatin modification analysis. Nucleic Acids Res., 41, 9230–9242.

5. Yang, X., Shao, X., Gao, L. and Zhang, S. (2015) Systematic DNA methylation analysis of multiple cell lines reveals common and specific patterns within and across tissues of origin. Hum. Mol. Genet., 24, 4374–4384.

6. Wang, C. and Zhang, S. (2017) Large-scale determination and characterization of cell type-specific regulatory elements in the human genome. J. Mol. Cell. Biol., 9, 463–476.

7. Butler, J.E. and Kadonaga, J.T. (2002) The RNA polymerase II core promoter: a key component in the regulation of gene expression. Genes Dev., 16, 2583–2592.

8. Leung, K.K., Ng, L.J., Ho, K.K., Tam, P.P. and Cheah, K.S. (1998) Different cis-regulatory DNA elements mediate developmental stage-and tissue-specific expression of the human COL2A1 gene in transgenic mice. J. Cell Biol, 141, 1291–1300.

9. Ye, Y., Gao, L. and Zhang, S. (2019) MSTD: an efficient method for detecting multiscale topological domains from symmetric and asymmetric 3D genomic maps. Nucleic Acids Res., 47, e65–e65.

10. Cairns, J., Freire-Pritchett, P., Wingett, S.W., Várnai, C., Dimond, A., Plagnol, V., Zerbino, D., Schoenfelder, S., Javierre, B.-M. and Osborne, C. (2016) CHiCAGO: robust detection of DNA looping interactions in Capture Hi-C data. Genome Biol., 17, 127.

11. Fullwood, M.J., Liu, M.H., Pan, Y.F., Liu, J., Xu, H., Mohamed, Y.B., Orlov, Y.L., Velkov, S., Ho, A. and Mei, P.H. (2009) An oestrogen-receptor-α-bound human chromatin interactome. Nature, 462, 58–64.

12. Lieberman-Aiden, E., Van Berkum, N.L., Williams, L., Imakaev, M., Ragoczy, T., Telling, A., Amit, I., Lajoie, B.R., Sabo, P.J. and Dorschner, M.O. (2009) Comprehensive mapping of long-range interactions reveals folding principles of the human genome. Science, 326, 289–293.

13. Schoenfelder, S., Furlan-Magaril, M., Mifsud, B., Tavares-Cadete, F., Sugar, R., Javierre, B.-M., Nagano, T., Katsman, Y., Sakthidevi, M. and Wingett, S.W. (2015) The pluripotent regulatory circuitry connecting promoters to their long-range interacting elements. Genome Res., 25, 582–597.

14. Corces, M.R., Granja, J.M., Shams, S., Louie, B.H., Seoane, J.A., Zhou, W., Silva, T.C., Groeneveld, C., Wong, C.K. and Cho, S.W. (2018) The chromatin accessibility landscape of primary human cancers. Science, 362, eaav1898.

15. Cao, Q., Anyansi, C., Hu, X., Xu, L., Xiong, L., Tang, W., Mok, M.T., Cheng, C., Fan, X. and Gerstein, M. (2017) Reconstruction of enhancer–target networks in 935 samples of human primary cells, tissues and cell lines. Nat. Genet., 49, 1428–1436.

16. Roy, S., Siahpirani, A.F., Chasman, D., Knaack, S., Ay, F., Stewart, R., Wilson, M. and Sridharan, R. (2015) A predictive modeling approach for cell line-specific long-range regulatory interactions. Nucleic Acids Res., 43, 8694–8712.

17. Whalen, S., Truty, R.M. and Pollard, K.S. (2016) Enhancer–promoter interactions are encoded by complex genomic signatures on looping chromatin. Nat. Genet., 48, 488–496.

18. Zhu, Y., Chen, Z., Zhang, K., Wang, M., Medovoy, D., Whitaker, J.W., Ding, B., Li, N., Zheng, L. and Wang, W. (2016) Constructing 3D interaction maps from 1D epigenomes. Nat. Commun., 7, 10812.

19. Buenrostro, J.D., Wu, B., Litzenburger, U.M., Ruff, D., Gonzales, M.L., Snyder, M.P., Chang, H.Y. and Greenleaf, W.J. (2015) Single-cell chromatin accessibility reveals principles of regulatory variation. Nature, 523, 486–490.

20. Cusanovich, D.A., Daza, R., Adey, A., Pliner, H.A., Christiansen, L., Gunderson, K.L., Steemers, F.J., Trapnell, C. and Shendure, J. (2015) Multiplex single-cell profiling of chromatin accessibility by combinatorial cellular indexing. Science, 348, 910–914.

21. Felsenfeld, G., Boyes, J., Chung, J., Clark, D. and Studitsky, V. (1996) Chromatin structure and gene expression. Proc. Natl. Acad. Sci. U. S. A., 93, 9384–9388.

22. Klemm, S.L., Shipony, Z. and Greenleaf, W.J. (2019) Chromatin accessibility and the regulatory epigenome. Nat. Rev. Genet., 20, 207–220.

23. Cusanovich, D.A., Hill, A.J., Aghamirzaie, D., Daza, R.M., Pliner, H.A., Berletch, J.B., Filippova, G.N., Huang, X., Christiansen, L. and DeWitt, W.S. (2018) A single-cell atlas of in vivo mammalian chromatin accessibility. Cell, 174, 1309–1324.

24. Cusanovich, D.A., Reddington, J.P., Garfield, D.A., Daza, R.M., Aghamirzaie, D., Marco-Ferreres, R., Pliner, H.A., Christiansen, L., Qiu, X. and Steemers, F.J. (2018) The cis-regulatory dynamics of embryonic development at single-cell resolution. Nature, 555, 538–542.

25. Clark, S.J., Lee, H.J., Smallwood, S.A., Kelsey, G. and Reik, W. (2016) Single-cell epigenomics: powerful new methods for understanding gene regulation and cell identity. Genome Biol., 17, 72.

26. Pliner, H.A., Packer, J.S., McFaline-Figueroa, J.L., Cusanovich, D.A., Daza, R.M., Aghamirzaie, D., Srivatsan, S., Qiu, X., Jackson, D. and Minkina, A. (2018) Cicero predicts cis-regulatory DNA Interactions from single-cell chromatin accessibility data. Mol. Cell, 71, 858–871.

27. Tung, P.-Y., Blischak, J.D., Hsiao, C.J., Knowles, D.A., Burnett, J.E., Pritchard, J.K. and Gilad, Y. (2017) Batch effects and the effective design of single-cell gene expression studies. Sci. Rep., 7, 39921.

28. Zhang, Y., Liu, T., Meyer, C.A., Eeckhoute, J., Johnson, D.S., Bernstein, B.E., Nusbaum, C., Myers, R.M., Brown, M. and Li, W. (2008) Model-based analysis of ChIP-Seq (MACS). Genome Biol., 9, R137.

29. Saunders, A., Macosko, E.Z., Wysoker, A., Goldman, M., Krienen, F.M., de Rivera, H., Bien, E., Baum, M., Bortolin, L. and Wang, S. (2018) Molecular diversity and specializations among the cells of the adult mouse brain. Cell, 174, 1015–1030.

30. Consortium, T.M. (2018) Single-cell transcriptomics of 20 mouse organs creates a Tabula Muris. Nature, 562, 367–372.

31. Han, X., Wang, R., Zhou, Y., Fei, L., Sun, H., Lai, S., Saadatpour, A., Zhou, Z., Chen, H. and Ye, F. (2018) Mapping the mouse cell atlas by microwell-seq. Cell, 172, 1091–1107.

32. Consortium, E.P. (2012) An integrated encyclopedia of DNA elements in the human genome. Nature, 489, 57–74.

33. van den Boogaard, M., Wong, L.E., Tessadori, F., Bakker, M.L., Dreizehnter, L.K., Wakker, V., Bezzina, C.R., AC’t Hoen, P., Bakkers, J. and Barnett, P. (2012) Genetic variation in T-box binding element functionally affects SCN5A/SCN10A enhancer. J Clin Invest, 122, 2519–2530.

34. Li, B., Qing, T., Zhu, J., Wen, Z., Yu, Y., Fukumura, R., Zheng, Y., Gondo, Y. and Shi, L. (2017) A comprehensive mouse transcriptomic BodyMap across 17 tissues by RNA-seq. Sci. Rep., 7, 4200.

35. Xiao, S.-J., Zhang, C., Zou, Q. and Ji, Z.-L. (2010) TiSGeD: a database for tissue-specific genes. Bioinformatics, 26, 1273–1275.

36. Breiman, L. (1996) Bagging predictors. Mach. Learn., 24, 123–140.

37. Maaten, L.v.d. and Hinton, G. (2008) Visualizing data using t-SNE. J. Mach. Learn. Res., 9, 2579–2605.

38. Beygelzimer, A., Kakadet, S., Langford, J., Arya, S., Mount, D. and Li, S. (2013) FNN: fast nearest neighbor search algorithms and applications. R package version, 1.

39. Danaher, P., Wang, P. and Witten, D.M. (2014) The joint graphical lasso for inverse covariance estimation across multiple classes. J. R. Stat. Soc. Ser. B-Stat. Methodol., 76, 373–397.

40. Friedman, J., Hastie, T. and Tibshirani, R. (2008) Sparse inverse covariance estimation with the graphical lasso. Biostatistics, 9, 432–441.

41. Yuan, M. and Lin, Y. (2007) Model selection and estimation in the Gaussian graphical model. Biometrika, 94, 19–35.

42. Javierre, B.M., Burren, O.S., Wilder, S.P., Kreuzhuber, R., Hill, S.M., Sewitz, S., Cairns, J., Wingett, S.W., Várnai, C. and Thiecke, M.J. (2016) Lineage-specific genome architecture links enhancers and non-coding disease variants to target gene promoters. Cell, 167, 1369–1384.

43. Haeussler, M., Zweig, A.S., Tyner, C., Speir, M.L., Rosenbloom, K.R., Raney, B.J., Lee, C.M., Lee, B.T., Hinrichs, A.S. and Gonzalez, J.N. (2018) The UCSC Genome Browser database: 2019 update. Nucleic Acids Res., 47, D853–D858.

44. Stuart, T., Butler, A., Hoffman, P., Hafemeister, C., Papalexi, E., Mauck III, W.M., Hao, Y., Stoeckius, M., Smibert, P. and Satija, R. (2019) Comprehensive Integration of Single-Cell Data. Cell, 177, 1888–1902.

45. Yu, G., Wang, L.-G., Han, Y. and He, Q.-Y. (2012) clusterProfiler: an R package for comparing biological themes among gene clusters. OMICS, 16, 284–287.

46. Heinz, S., Benner, C., Spann, N., Bertolino, E., Lin, Y.C., Laslo, P., Cheng, J.X., Murre, C., Singh, H. and Glass, C.K. (2010) Simple combinations of lineage-determining transcription factors prime cis-regulatory elements required for macrophage and B cell identities. Mol. Cell, 38, 576–589.

47. Yoshida, H., Lareau, C.A., Ramirez, R.N., Rose, S.A., Maier, B., Wroblewska, A., Desland, F., Chudnovskiy, A., Mortha, A. and Dominguez, C. (2019) The cis-Regulatory Atlas of the Mouse Immune System. Cell, 176, 897–912.

48. Lacorazza, H.D., Miyazaki, Y., Di Cristofano, A., Deblasio, A., Hedvat, C., Zhang, J., Cordon-Cardo, C., Mao, S., Pandolfi, P.P. and Nimer, S.D. (2002) The ETS protein MEF plays a critical role in perforin gene expression and the development of natural killer and NK-T cells. Immunity, 17, 437–449.

49. Wasylyk, C., Schlumberger, S.E., Criqui-Filipe, P. and Wasylyk, B. (2002) Sp100 interacts with ETS-1 and stimulates its transcriptional activity. Mol. Cell. Biol., 22, 2687–2702.

50. Fu, J., Gaetani, S., Oveisi, F., Verme, J.L., Serrano, A., de Fonseca, F.R., Rosengarth, A., Luecke, H., Di Giacomo, B. and Tarzia, G. (2003) Oleylethanolamide regulates feeding and body weight through activation of the nuclear receptor PPAR-α. Nature, 425, 90–93.

51. Yue, F., Cheng, Y., Breschi, A., Vierstra, J., Wu, W., Ryba, T., Sandstrom, R., Ma, Z., Davis, C. and Pope, B.D. (2014) A comparative encyclopedia of DNA elements in the mouse genome. Nature, 515, 355–364.

52. Seo, S., Lim, J.W., Yellajoshyula, D., Chang, L.W. and Kroll, K.L. (2007) Neurogenin and NeuroD direct transcriptional targets and their regulatory enhancers. EMBO J., 26, 5093–5108.

53. Li, Q.Y., Newbury-Ecob, R.A., Terrett, J.A., Wilson, D.I., Curtis, A.R., Yi, C.H., Gebuhr, T., Bullen, P.J., Robson, S.C. and Strachan, T. (1997) Holt-Oram syndrome is caused by mutations in TBX5, a member of the Brachyury (T) gene family. Nat. Genet., 15, 21-29.

54. Hoogaars, W.M., Engel, A., Brons, J.F., Verkerk, A.O., de Lange, F.J., Wong, L.E., Bakker, M.L., Clout, D.E., Wakker, V. and Barnett, P. (2007) Tbx3 controls the sinoatrial node gene program and imposes pacemaker function on the atria. Genes Dev., 21, 1098–1112.

55. Creyghton, M.P., Cheng, A.W., Welstead, G.G., Kooistra, T., Carey, B.W., Steine, E.J., Hanna, J., Lodato, M.A., Frampton, G.M. and Sharp, P.A. (2010) Histone H3K27ac separates active from poised enhancers and predicts developmental state. Proc. Natl. Acad. Sci. U. S. A., 107, 21931–21936.

56. Arnolds, D.E., Liu, F., Fahrenbach, J.P., Kim, G.H., Schillinger, K.J., Smemo, S., McNally, E.M., Nobrega, M.A., Patel, V.V. and Moskowitz, I.P. (2012) TBX5 drives Scn5a expression to regulate cardiac conduction system function. J Clin Invest, 122, 2509–2518.

57. van den Boogaard, M., Smemo, S., Burnicka-Turek, O., Arnolds, D.E., van de Werken, H.J., Klous, P., McKean, D., Muehlschlegel, J.D., Moosmann, J. and Toka, O. (2014) A common genetic variant within SCN10A modulates cardiac SCN5A expression. J Clin Invest, 124, 1844–1852.

58. Doi, R., Oishi, K. and Ishida, N. (2010) CLOCK regulates circadian rhythms of hepatic glycogen synthesis through transcriptional activation of Gys2. J. Biol. Chem., 285, 22114–22121.

59. Mermet, J., Yeung, J., Hurni, C., Mauvoisin, D., Gustafson, K., Jouffe, C., Nicolas, D., Emmenegger, Y., Gobet, C. and Franken, P. (2018) Clock-dependent chromatin topology modulates circadian transcription and behavior. Genes Dev., 32, 347–358.

60. Klein, F.A., Pakozdi, T., Anders, S., Ghavi-Helm, Y., Furlong, E.E. and Huber, W. (2015) FourCSeq: analysis of 4C sequencing data. Bioinformatics, 31, 3085–3091.

61. Huang, J., Elicker, J., Bowens, N., Liu, X., Cheng, L., Cappola, T.P., Zhu, X. and Parmacek, M.S. (2012) Myocardin regulates BMP10 expression and is required for heart development. J Clin Invest, 122, 3678–3691.

62. Chauveau, C., Bonnemann, C.G., Julien, C., Kho, A.L., Marks, H., Talim, B., Maury, P., Arne-Bes, M.C., Uro-Coste, E. and Alexandrovich, A. (2013) Recessive TTN truncating mutations define novel forms of core myopathy with heart disease. Hum. Mol. Genet., 23, 980–991.

63. Granados-Riveron, J.T., Ghosh, T.K., Pope, M., Bu’Lock, F., Thornborough, C., Eason, J., Kirk, E.P., Fatkin, D., Feneley, M.P. and Harvey, R.P. (2010) α-Cardiac myosin heavy chain (MYH6) mutations affecting myofibril formation are associated with congenital heart defects. Hum. Mol. Genet., 19, 4007–4016.

64. Yoon, J.-H., Abdelmohsen, K., Srikantan, S., Yang, X., Martindale, J.L., De, S., Huarte, M., Zhan, M., Becker, K.G. and Gorospe, M. (2012) LincRNA-p21 suppresses target mRNA translation. Mol. Cell, 47, 648–655.

65. Zhi, H., Ning, S., Li, X., Li, Y., Wu, W. and Li, X. (2014) A novel reannotation strategy for dissecting DNA methylation patterns of human long intergenic non-coding RNAs in cancers. Nucleic Acids Res., 42, 8258–8270.

66. Corces, M.R., Buenrostro, J.D., Wu, B., Greenside, P.G., Chan, S.M., Koenig, J.L., Snyder, M.P., Pritchard, J.K., Kundaje, A. and Greenleaf, W.J. (2016) Lineage-specific and single-cell chromatin accessibility charts human hematopoiesis and leukemia evolution. Nat. Genet., 48, 1193–1203.

67. Preissl, S., Fang, R., Huang, H., Zhao, Y., Raviram, R., Gorkin, D.U., Zhang, Y., Sos, B.C., Afzal, V. and Dickel, D.E. (2018) Single-nucleus analysis of accessible chromatin in developing mouse forebrain reveals cell-type-specific transcriptional regulation. Nat. Neurosci., 21, 432–439.

68. Trapnell, C., Cacchiarelli, D., Grimsby, J., Pokharel, P., Li, S., Morse, M., Lennon, N.J., Livak, K.J., Mikkelsen, T.S. and Rinn, J.L. (2014) The dynamics and regulators of cell fate decisions are revealed by pseudotemporal ordering of single cells. Nat. Biotechnol., 32, 381–386.

