## Supplementary Materials for "Joint reconstruction of *cis*-regulatory interaction networks across multiple tissues using single-cell chromatin accessibility data"

### Supplementary Figures

**Figure S1.** Cell number and data sparsity of single-cell ATAC-seq data for each tissue.
**Figure S2.** Illustration of the workflow of data processing for each tissue.
**Figure S3.** Number of interacting co-accessible DNA element pairs in each tissue estimated by JRIM and by Cicero.
**Figure S4.** Distribution of distance between interacting co-accessible DNA elements.
**Figure S5.** Enrichment degrees (fold changes) of the number of *cis*-regulatory interactions in different genomic regions.
**Figure S6.** Mean number of occurring tissues of six types of genomic region related interactions.
**Figure S7.** Comparison of the number of promoter-related interactions of 27 consistently and highly expressed genes and remaining genes.
**Figure S8.** Hierarchical clustering of the 13 tissues in terms of gene activity scores.
**Figure S9.** Enrichment analysis of chromatin modification mark around TSSs of tissue-specific differential activity genes.
**Figure 10.** The H3K4me1 signal and CTCF signal of tissue-specific functional peaks compared to those of other peaks.
**Figure S11.** Reconstructed regulatory networks around *Fto* gene and *Irx3* gene.
**Figure S12.** Illustration of 4C-seq data of *Gys2* gene in liver at different time points in wild-type mouse and clock-deficient Bmal1 knockout mouse.
**Figure S13.** Spatial regulatory loci interacting with *Gys2* TSS in liver and kidney estimated by 'FourCSeq' method from a 4C-seq data.
**Figure S14.** Changes of sparsity and similarity of *cis*-regulatory interaction networks with respect to parameter tuning.

### Supplementary Tables

**Table S1.** Examples of Homer motif analysis for tissue-specific functional peaks.
**Table S2.** Lists of differential activity genes (sorted by z-score).
**Table S3.** Related genes of common interactions among immune and nervous tissues respectively.
**Table S4.** GO terms enriched in differential activity genes.
**Table S5.** GO terms enriched in related genes of common interactions among immune tissues.
**Table S6.** GO terms enriched in related genes of common interactions among nervous tissues.

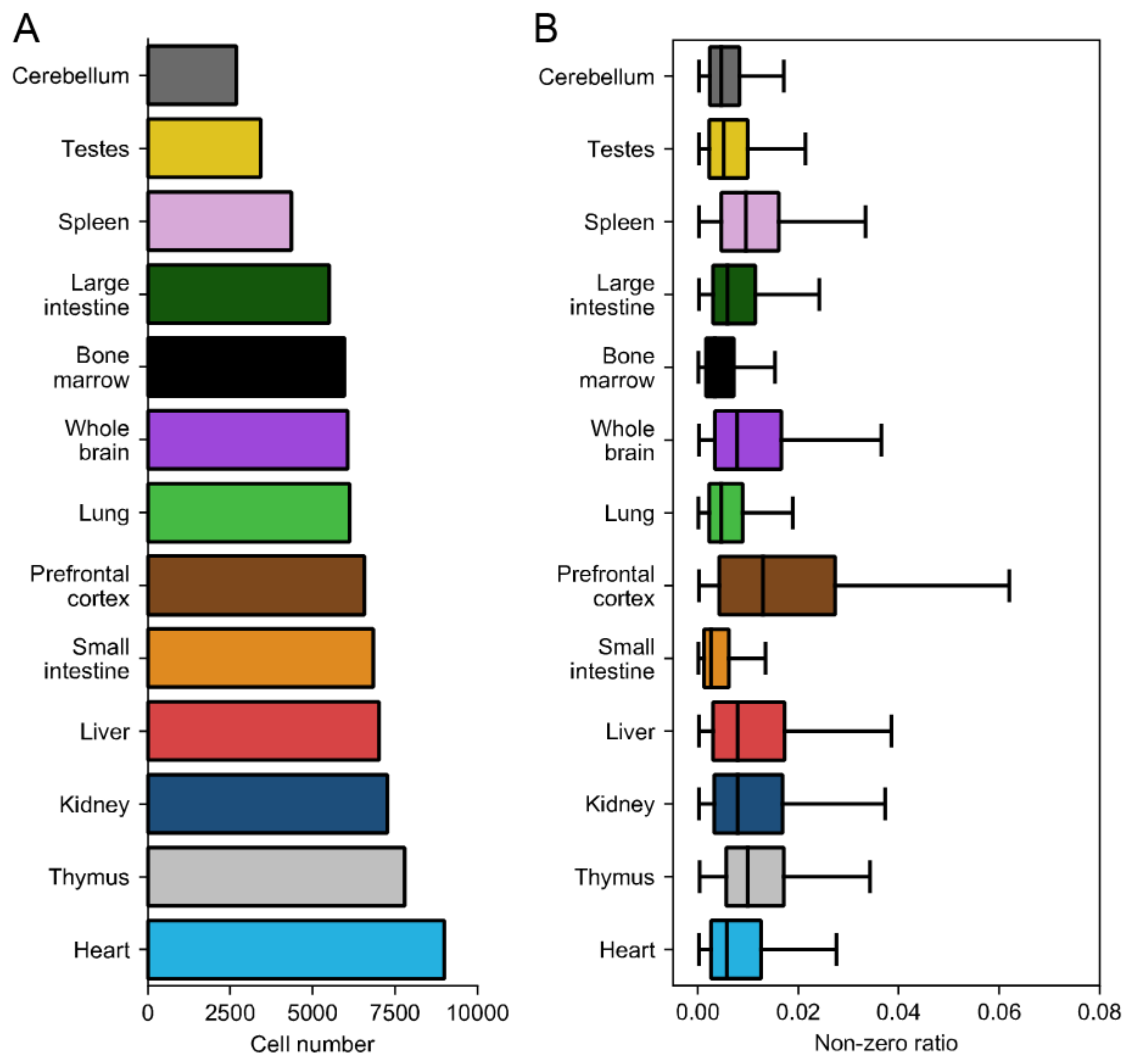


**Figure S1.** Cell number (A) and data sparsity (B) of single-cell ATAC-seq data for each tissue.

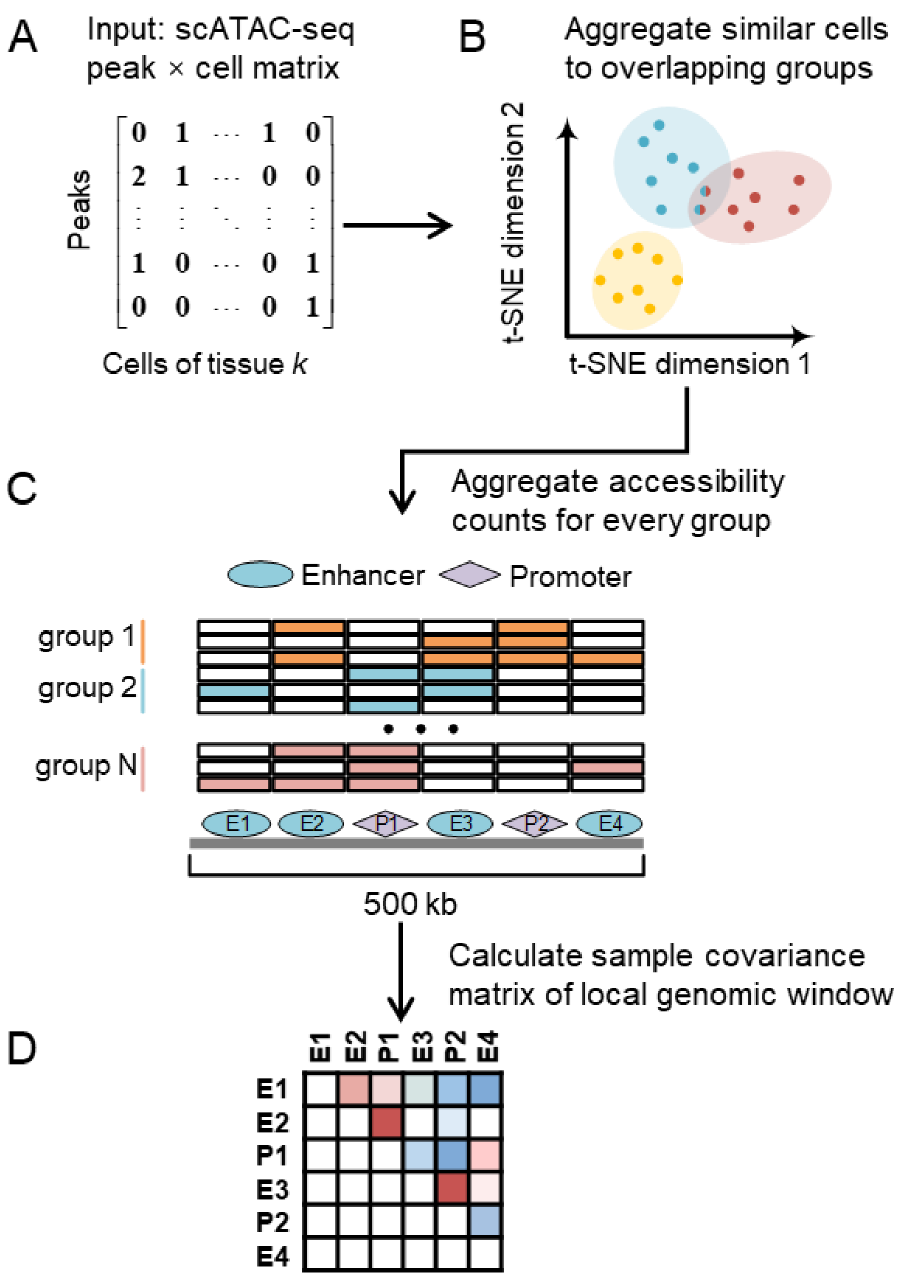


**Figure S2.** Illustration of the workflow of data processing for each tissue. (A) JRIM takes the peak × cell matrix of each tissue as input. (B) JRIM first maps single cells into low dimensional spaces using t-SNE and aggregates single cells into overlapping groups. (C) After clustering similar cells into overlapping groups, JRIM aggregates their accessibility counts to construct grouped matrix. (D) Finally, JRIM calculates sample covariance matrix of grouped matrix for each local genomic window. The resulting sample covariance matrices are adopted to joint graphical lasso model to jointly reconstruct *cis*-regulatory interaction networks (Figure 1).

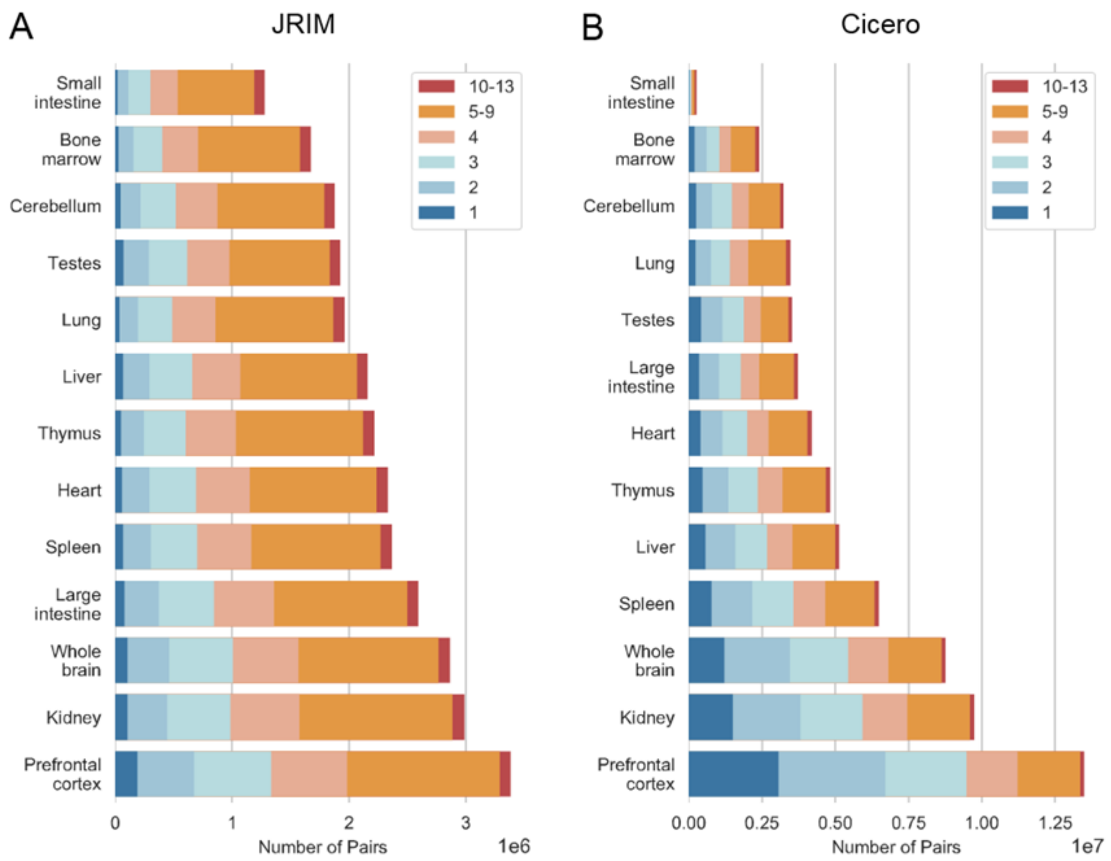


**Figure S3.** Number of interacting co-accessible DNA element pairs (i.e., *cis*-regulatory interactions) in each tissue estimated by JRIM (A), and by Cicero (using the same sparsity parameter λ=0.25 for each tissue respectively) (B).

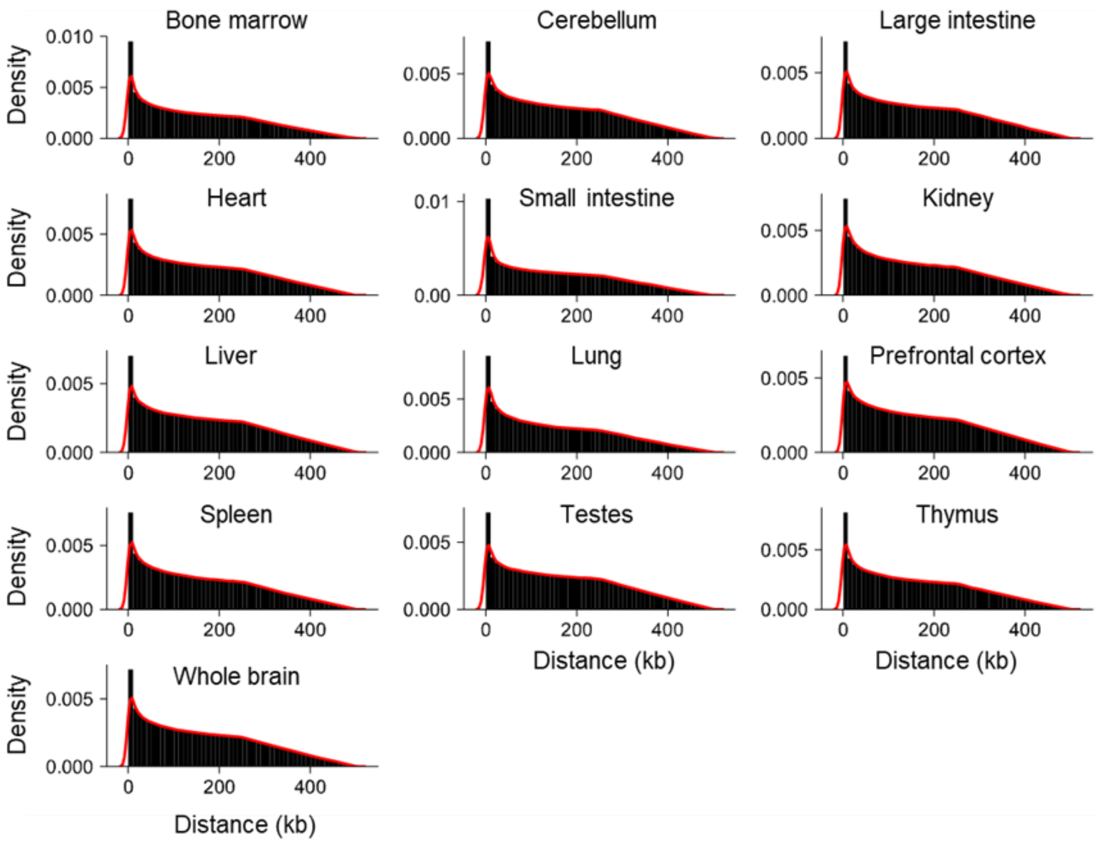


**Figure S4.** Distribution of distance between interacting co-accessible DNA elements.

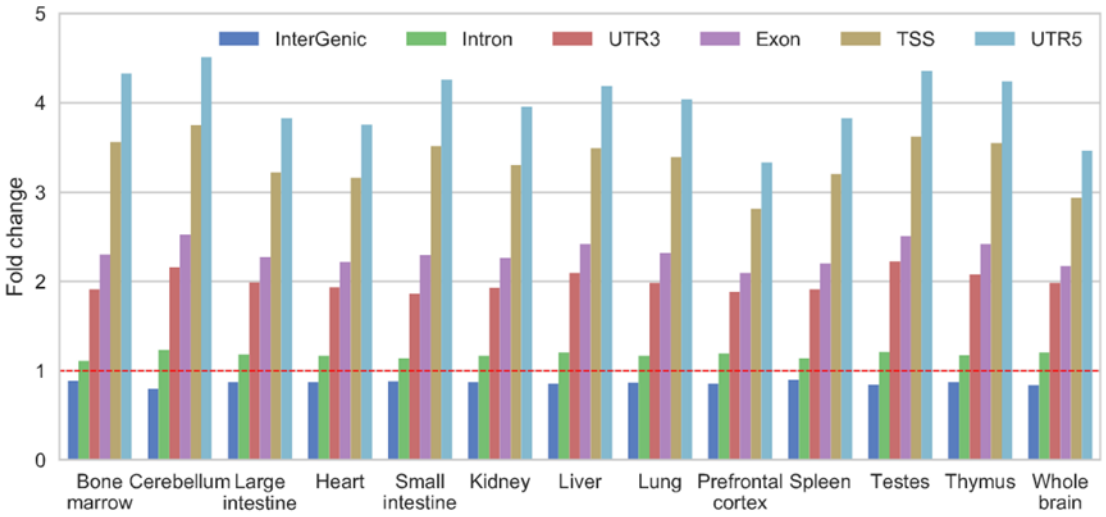


**Figure S5.** Enrichment degrees (fold changes) of the number of *cis*-regulatory interactions in different genomic regions.

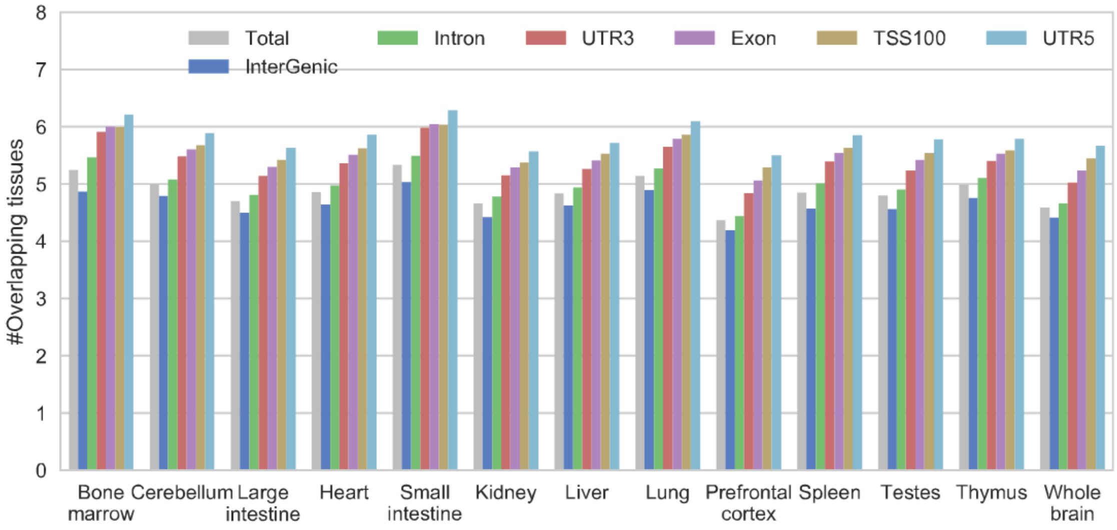


**Figure S6.** Mean number of occurring tissues of six types of genomic region related interactions.

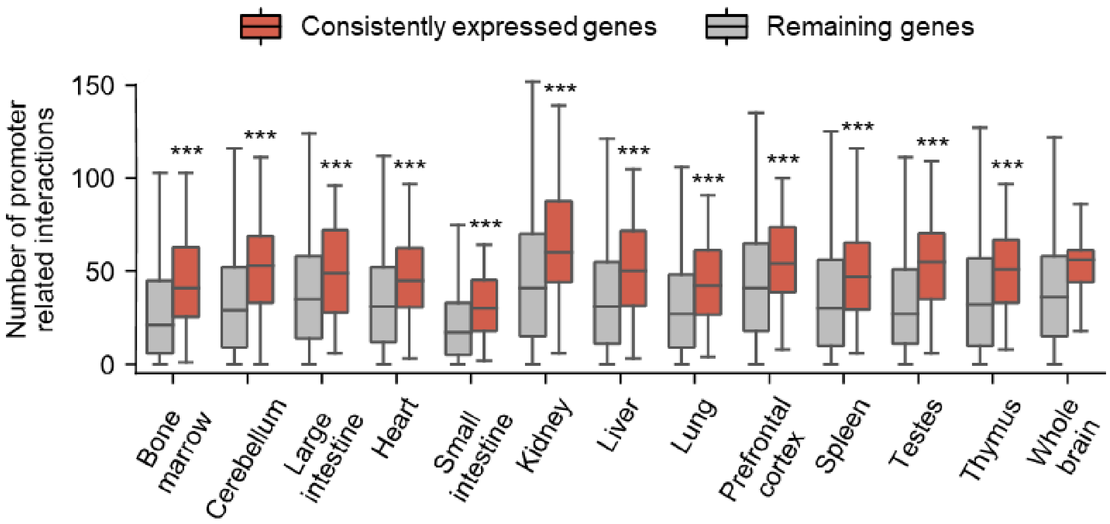


**Figure S7.** Comparison of the number of promoter-related interactions of 27 consistently and highly expressed genes obtained from (1) and remaining genes. Statistical significance of the difference was calculated using two-sample Wilcoxon tests with $P < 0.01$ in all 13 tissues.

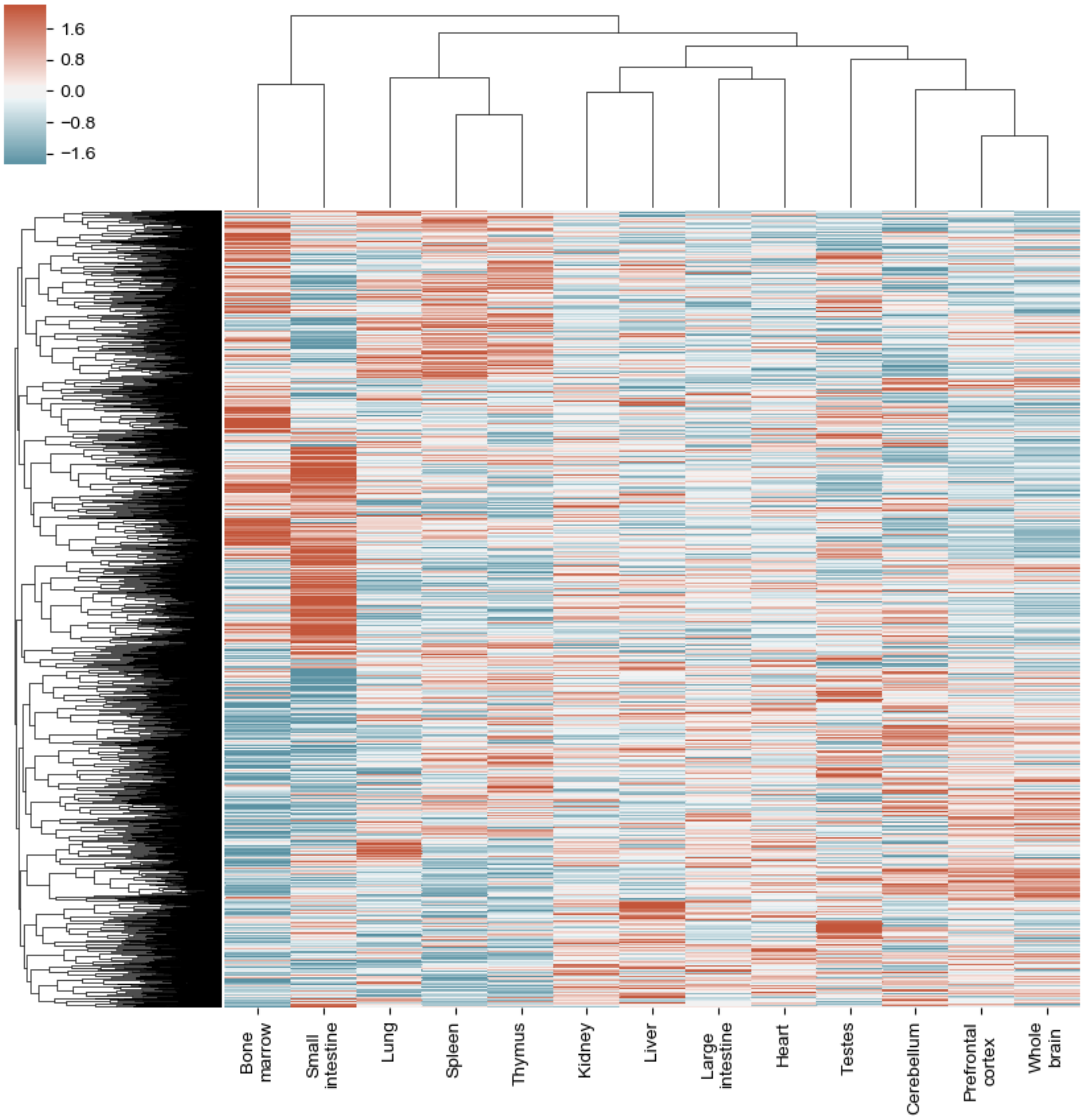


**Figure S8.** Hierarchical clustering of the 13 tissues in terms of gene activity scores.

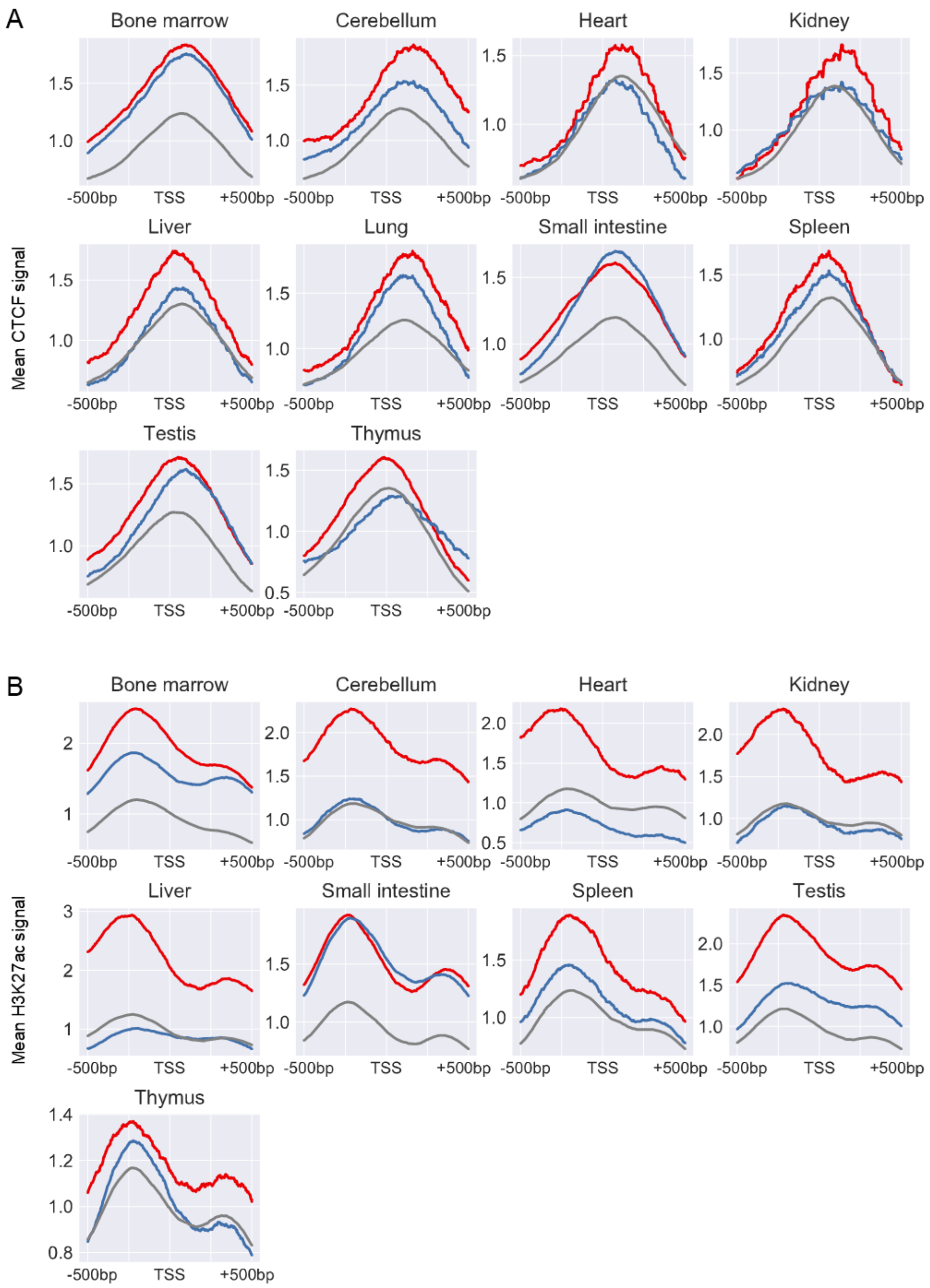


**Figure S9.** Enrichment analysis of chromatin modification mark around TSSs of tissue-specific differential activity genes (DAGs). (A) Enrichment analysis of CTCF around TSSs of DAGs. The CTCF signal around TSSs of DAGs and remaining genes are labeled as red and gray colors, respectively. The blue line is the mean CTCF signals around TSS of DAGs in other tissues. (B) Enrichment analysis of H3K27ac mark around TSSs of DAGs. The colors of lines are the same as (A).

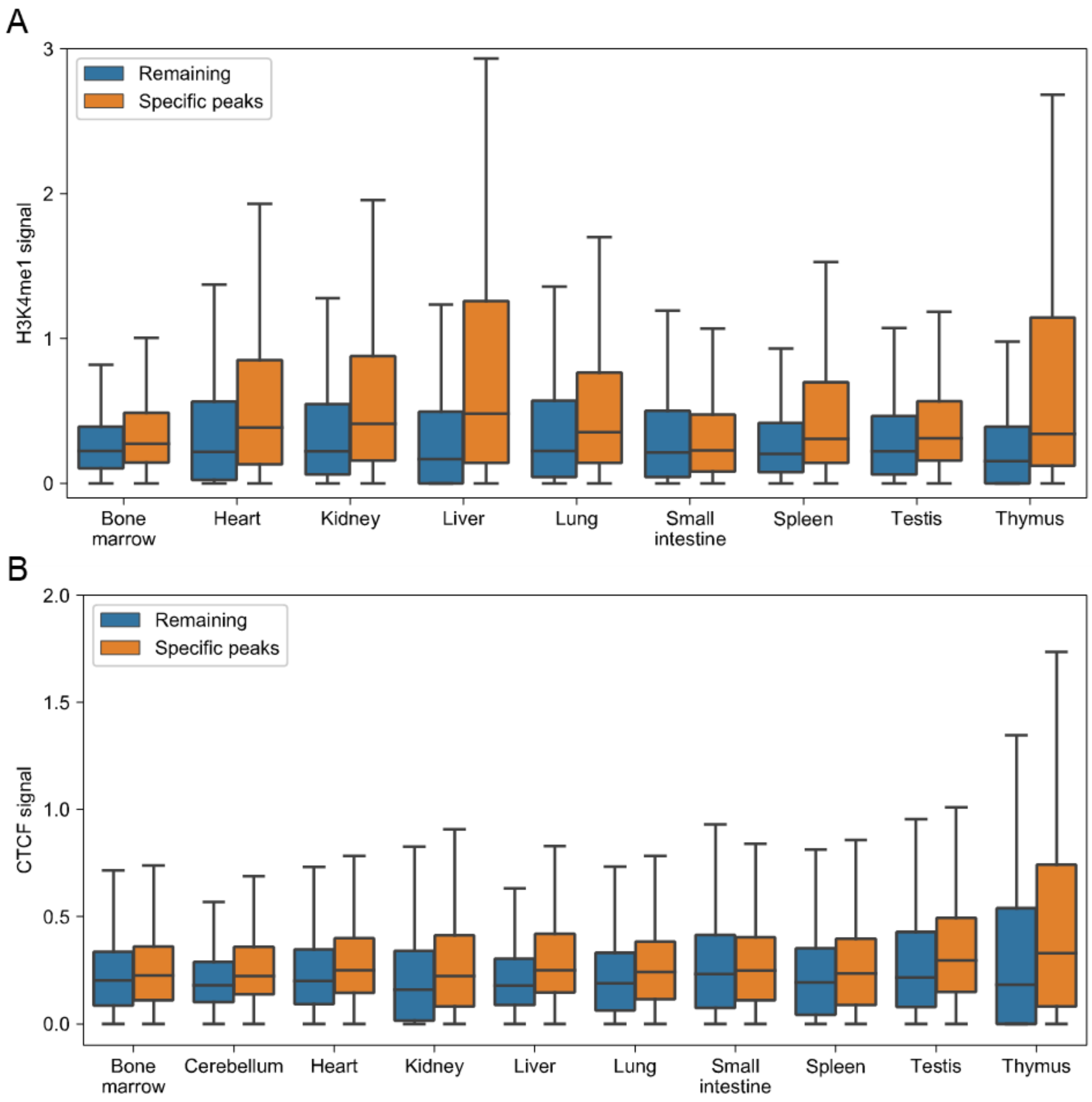


**Figure 10.** The H3K4me1 signal (A) and CTCF signal (B) of tissue-specific functional peaks compared to those of other peaks.

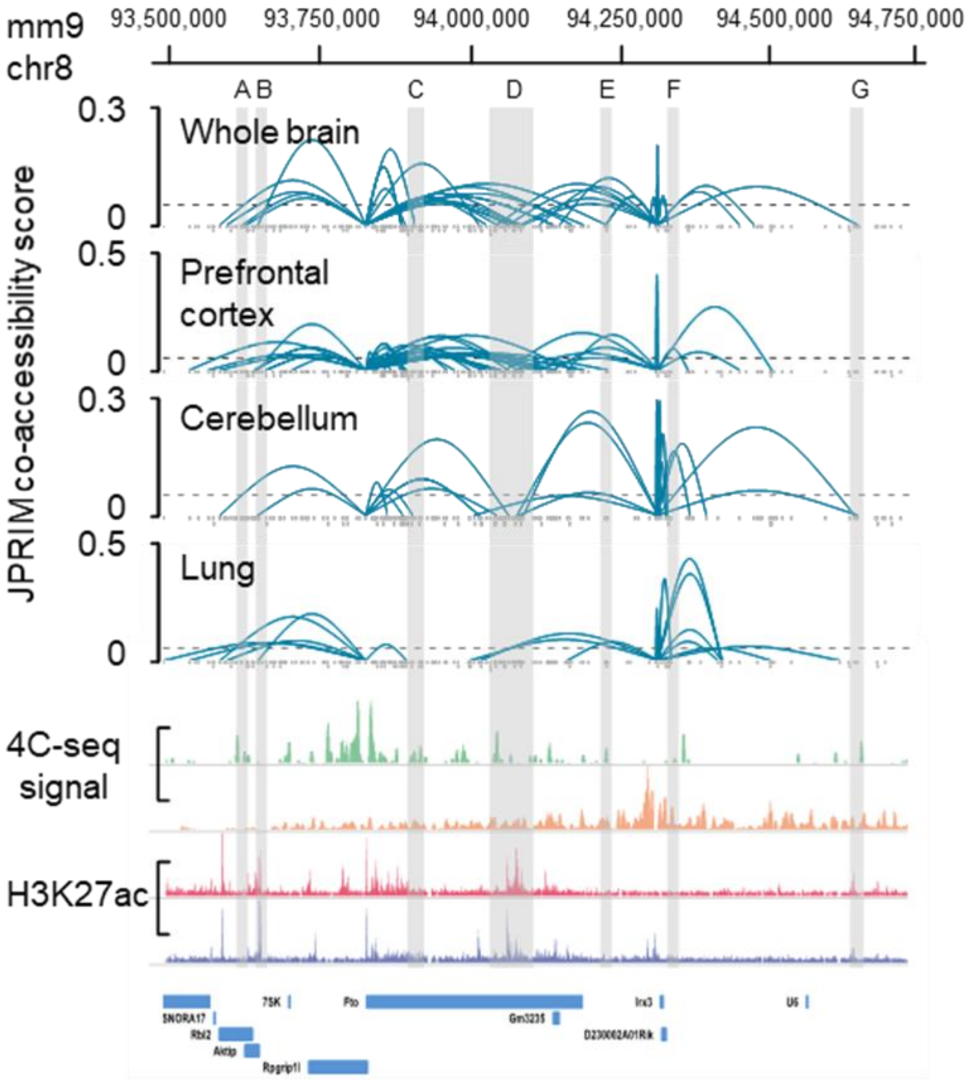


**Figure S11.** Reconstructed regulatory networks around *Fto* gene and *Irx3* gene (chr9: 93,500,000-94,750,000). The green line and yellow line represent the contact frequencies with *Fto* TSS and *Irx3* TSS respectively. The red line and blue line are the ChIP-seq signal of H3K27ac in cortex and cerebellum respectively. *Irx3* is highly expressed in brain and lung. *Fto* is highly expressed in brain only. JRIM identifies the relatively high activity of *Fto* in brain and prefrontal cortex. And it has been reported (2) that enhancers located in *Fto* region regulate the expression of *Irx3* in brain as shown in the region D.

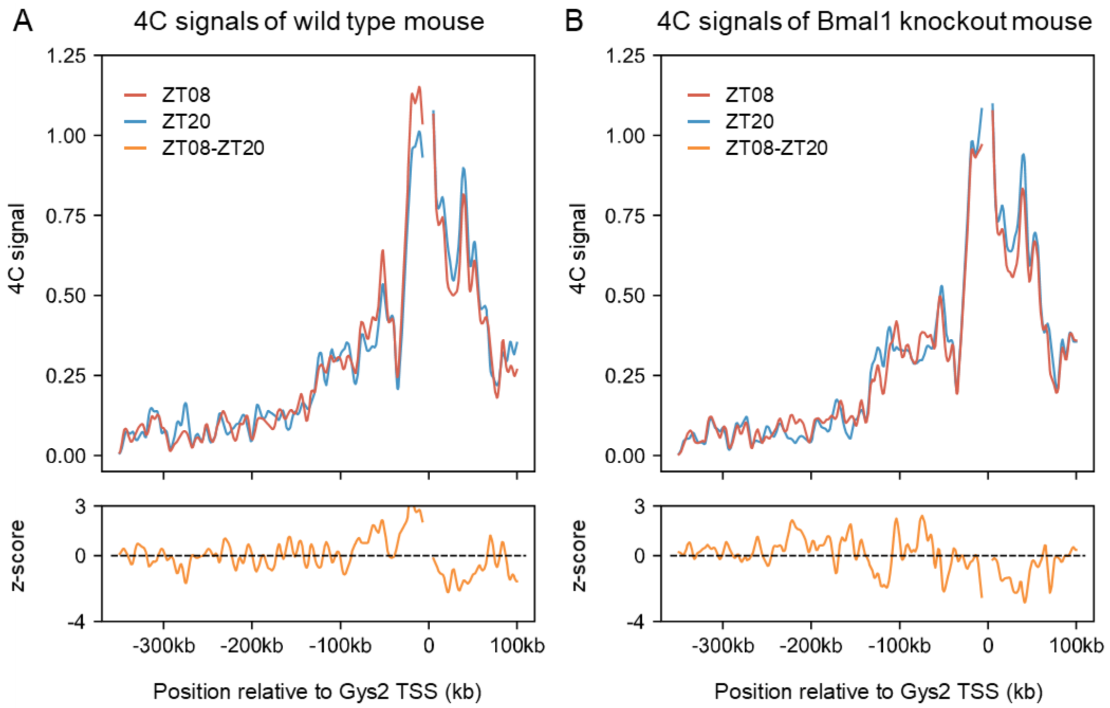


**Figure S12.** Illustration of 4C-seq data of Gys2 gene in liver at different time points in wild-type mouse (A) and clock-deficient Bmal1 knockout mouse (B). 'ZT08' and 'ZT20' indicate that the 4C-seq experiment is performed at zeitgeber time 8 and zeitgeber time 20. 'ZT08-ZT20' indicates the difference between signals at ZT08 and ZT20.

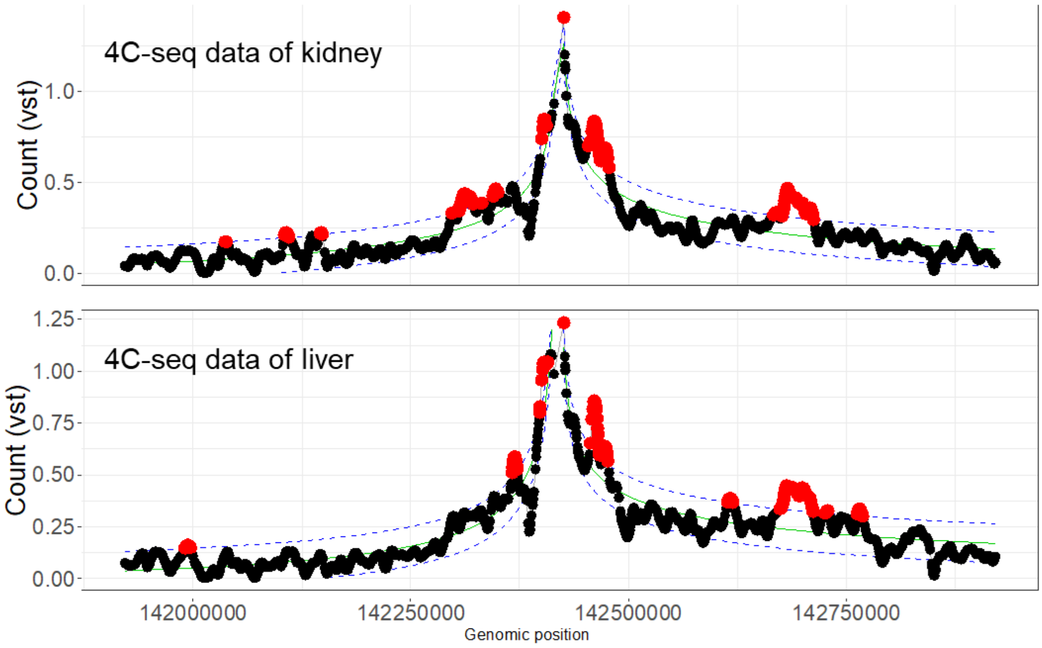


**Figure S13.** Spatial regulatory loci interacting with *Gys2* TSS in liver and kidney estimated by 'FourCSeq' method (3) from a 4C-seq data with z-score >1.5.

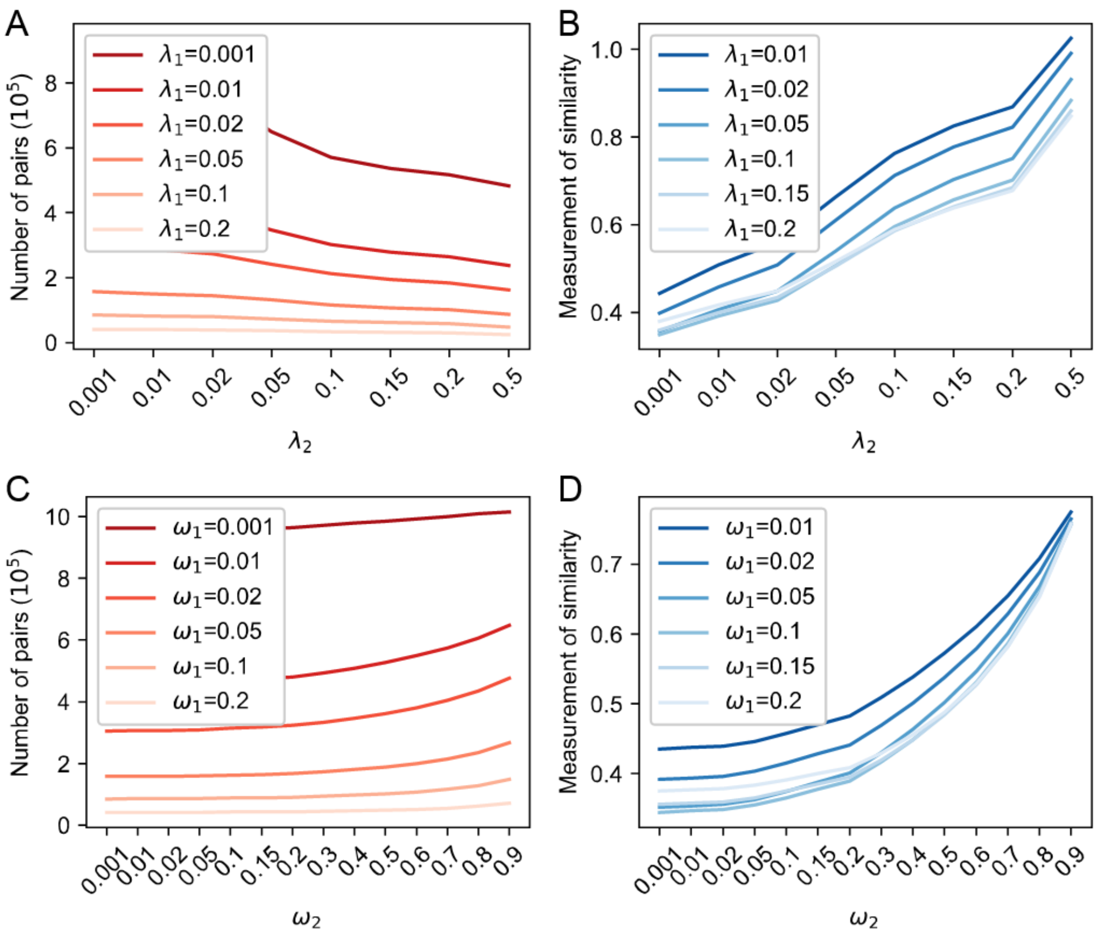


**Figure S14.** Changes of sparsity and similarity of *cis*-regulatory interaction networks with respect to parameter tuning: $\lambda_1$ and $\lambda_2$ in (A, B) and $\omega_1$ and $\omega_2$ in (C, D). This experiment was performed in four tissues (kidney, liver, heart and thymus).

**Table S1.** Examples of Homer motif analysis for tissue-specific functional peaks. Top three enriched motifs of each tissue are shown.

| Tissue | Motif | *P*-value | TF | Brief description | Reference |
|---|---|---|---|---|---|
| Bone marrow | 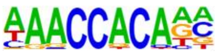 | 1e-45 | RUNX1 | RUNX1 is essential for the development of normal hematopoiesis and Involved in lineage commitment of immature T cell precursors. | 18258917 |
| | 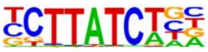 | 1e-44 | GATA6 | | |
| | 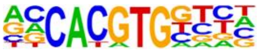 | 1e-43 | MAX | MAX is overexpressed in peripheral blood mononuclear cells, CD4 T cells, and monocytes. | 17072327 |
| Cerebellum | 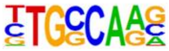 | 1e-105 | NF1 | | |
| | 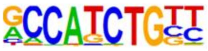 | 1e-100 | NEURO D1 | NEUROD1 associates with chromatin to enhancer regulatory elements in neurogenesis regulation genes. | 18007592 |
| | 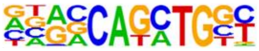 | 1e-99 | ATOH1 | ATOH1 plays a role in the differentiation of subsets of neural cells by activating E box-dependent transcription. | 10648228 |
| Large intestine | 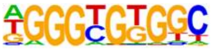 | 1e-229 | KLF5 | Tissue-specific expression in digestive track. Highest expression in adult mouse colon. | 25409824 |
| | 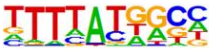 | 1e-217 | HOXA11 | | |
| | 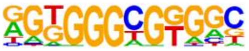 | 1e-211 | KLF14 | | |
| Heart | 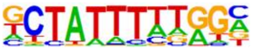 | 1e-161 | MEF2B | MEF2B may be involved in muscle-specific and growth factor-related transcription. | 9443808 |
| | 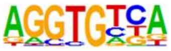 | 1e-153 | TBX5 | TBX5 regulates the transcription of ion channel genes and is essential for heart development. | 20133910 |
| | 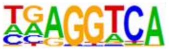 | 1e-149 | THRB | | |
| Small intestine | 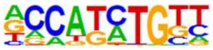 | 1e-77 | OLIG2 | | |
| | 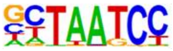 | 1e-67 | CRX | | |
| | 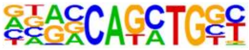 | 1e-65 | ATOH1 | Express specificity in adult epithelial cells of the gastrointestinal tract. | |
| Kidney | 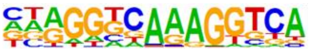 | 1e-282 | PPARA | PPARA is key regulator of lipid metabolism and regulates the peroxisomal beta-oxidation pathway of fatty acids. | 12955147 |
| | 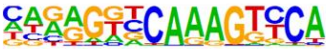 | 1e-277 | HNF4A | HNF4A binds fatty acids and may be essential for development of the liver, kidney and intestine. | 25409824 |
| | 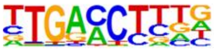 | 1e-248 | RARA | | |
| Liver | 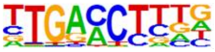 | 1e-313 | RARA | | |
| | 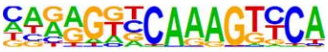 | 1e-283 | HNF4A | HNF4A binds fatty acids and may be essential for development of the liver, kidney and intestine. | 25409824 |
| | 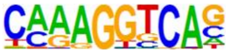 | 1e-272 | ERRA | | |
| Lung | 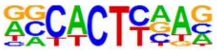 | 1e-119 | NKX2.1 | NKX2.1 forms a regulatory loop with GRHL2 that coordinates lung epithelial cell morphogenesis and differentiation. | 22955271 |
| | 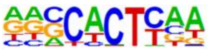 | 1e-114 | NKX2.5 | | |
| | 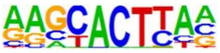 | 1e-108 | NKX3.1 | | |

| Tissue | Motif | *P*-value | TF | Brief description | Reference |
|---|---|---|---|---|---|
| Prefrontal cortex | 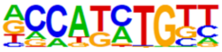 | 1e-156 | OLIG2 | OLIG2 is required for oligodendrocyte and motor neuron specification in the spinal cord. | 11955448 |
| | 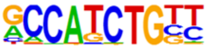 | 1e-143 | NEURO D1 | NEUROD1 associates with chromatin to enhancer regulatory elements in neurogenesis regulation genes. | 18007592 |
| | 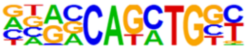 | 1e-123 | ATOH1 | ATOH1 plays a role in the differentiation of subsets of neural cells by activating E box-dependent transcription. | 10648228 |
| Spleen | 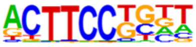 | 1e-136 | ELF4 | ELF4 plays a role in the development and function of NK and NK T-cells. | 12387738 |
| | 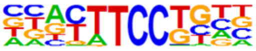 | 1e-128 | ETV2 | | |
| | 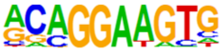 | 1e-123 | ETS1 | ETS1 controls the differentiation, survival and proliferation of lymphoid cells. | 11909962 |
| Testes | 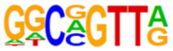 | 1e-102 | MYB | MYB plays an important role in the control of proliferation and differentiation of hematopoietic progenitor cells. | 20484083 |
| | 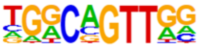 | 1e-88 | AMYB | AMYB acts as a master regulator of male meiosis by promoting expression of piRNAs. | 21750041 |
| | 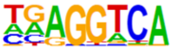 | 1e-85 | THRB | | |
| Thymus | 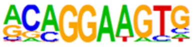 | 1e-247 | ETS1 | ETS1 controls the differentiation, survival and proliferation of lymphoid cells. | 11909962 |
| | 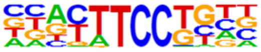 | 1e-226 | ETV2 | | |
| | 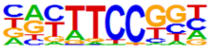 | 1e-213 | FLI1 | FLI1 is involved in erythroleukemia induction by Friend murine leukemia virus (F-MULV). | 2044959 |
| Whole brain | 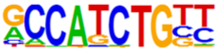 | 1e-164 | NEURO D1 | NEUROD1 associates with chromatin to enhancer regulatory elements in neurogenesis regulation genes. | 18007592 |
| | 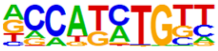 | 1e-163 | OLIG2 | OLIG2 is required for oligodendrocyte and motor neuron specification in the spinal cord. | 11955448 |
| | 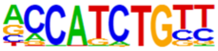 | 1e-151 | NEURO G2 | NEUROG2 is Involved in neuronal differentiation. | 14697366 |

### REFERENCES


1. Li, B., Qing, T., Zhu, J., Wen, Z., Yu, Y., Fukumura, R., Zheng, Y., Gondo, Y. and Shi, L. (2017) A comprehensive mouse transcriptomic BodyMap across 17 tissues by RNA-seq. *Sci. Rep.*, **7**, 4200.
2. Smemo, S., Tena, J.J., Kim, K.-H., Gamazon, E.R., Sakabe, N.J., Gómez-Marín, C., Aneas, I., Credidio, F.L., Sobreira, D.R. and Wasserman, N.F. (2014) Obesity-associated variants within FTO form long-range functional connections with IRX3. *Nature*, **507**, 371–375.
3. Klein, F.A., Pakozdi, T., Anders, S., Ghavi-Helm, Y., Furlong, E.E. and Huber, W. (2015) FourCSeq: analysis of 4C sequencing data. *Bioinformatics*, **31**, 3085-3091.
